# Small Molecule Approach to RNA Targeting Binder Discovery (SMARTBind) Using Deep Learning Without Structural Input

**DOI:** 10.1101/2025.09.24.678312

**Authors:** Shiyu Jiang, Amirhossein Taghavi, Tenghui Wang, Samantha M. Meyer, Jessica L. Childs-Disney, Chenglong Li, Matthew D. Disney, Yanjun Li

## Abstract

Accurate identification of small molecule binders to RNA is critical for chemical probes and therapeutics. Computational approaches offer a cost-effective strategy to identify small molecules targeting RNA but are often limited by poor predictive accuracy and high computational demands. Here, we introduce **<u>S</u>**mall **<u>M</u>**olecule **<u>A</u>**pproaches to **<u>R</u>**NA **<u>T</u>**argeting **Bind**er Discovery (SMARTBind), a structure-agnostic ligand discovery framework that combines an RNA large language model with contrastive learning and a ligand-specific decoy enhancement strategy. The RNA language model, pre-trained on millions of RNA sequences, together with the decoy enhancement strategy, addresses data scarcity and improves model generalizability. SMARTBind uses only RNA primary sequence to identify small molecule binders and their binding sites accurately. Across multiple benchmarks and case studies, SMARTBind outperforms existing data-driven and docking-based methods while significantly reducing computational cost. In a real-world application, SMARTBind identified novel small molecules targeting the precursor of oncogenic microRNA-21, validated by in vitro and cellular assays. These results highlight SMARTBind’s potential as a scalable, accurate, and structure-independent platform for RNA-targeted small molecule discovery.

## INTRODUCTION

RNAs are central regulators of cellular function and disease, and thus attractive therapeutic targets^1, 2^. Small molecules offer key advantages for RNA targeting, including oral bioavailability, brain penetration, and medicinal chemistry optimization^3^. However, identifying RNA-binding small molecules remains difficult, largely due to challenges in predicting RNA–ligand interactions accurately and efficiently. Traditional virtual screening relies on docking and scoring methods^4, 5, 6^, which have improved with machine learning^7, 8, 9, 10, 11^ but still struggle to recapitulate experimental results^12, 13^. Most tools are optimized for proteins and fail to capture RNA-specific features like high charge density, conformational flexibility, and limited structural data^14, 15, 16^. Existing RNA docking methods—including MORDOR^17^, Dock 6^18^, AutoDock^19^, LigandRNA^20^, and AnnapuRNA^21^—are limited by suboptimally performing scoring functions^22^ and are computationally intensive^16, 23^. To address these limitations, we developed SMARTBind (Small Molecule Approaches to RNA Targeting Binder Discovery), a deep learning framework that predicts RNA–ligand interactions and binding sites using only RNA sequence. SMARTBind leverages an RNA language model trained on millions of sequences to capture contextual and structural features and uses contrastive learning with a ligand-specific decoy enhancement strategy to build a unified RNA–ligand embedding space. This enables fast and accurate virtual screening without structural input and molecular docking process.

SMARTBind outperforms existing docking-based and data-driven methods in binder discovery across benchmarks and generalizes to diverse RNA targets regardless of conformational state or length. It scales efficiently—screening 10 billion compounds in 0.23 years on a single CPU core, compared to 3,000 years by docking^24^. In a real-world case, SMARTBind identified compounds targeting the precursor of oncogenic microRNA- 21 that inhibit biogenesis and induce apoptosis in triple-negative breast cancer cells. These findings demonstrate SMARTBind’s potential as a robust, structure-independent platform for RNA-targeted small molecule drug discovery.

## RESULTS

### Model overview

SMARTBind is a bi-functional RNA-targeted ligand discovery approach that estimates both binding score (interaction potential) and binding site (the region interacting with the active binder) using only the RNA’s sequence. As illustrated in Fig. 1a, given a target RNA sequence and a chemical library, SMARTBind identifies promising binders by ranking candidates based on predicted binding scores. Its binding site prediction module then suggests likely RNA interaction regions for these hits, providing insights for binding mechanisms.

**Fig. 1.**
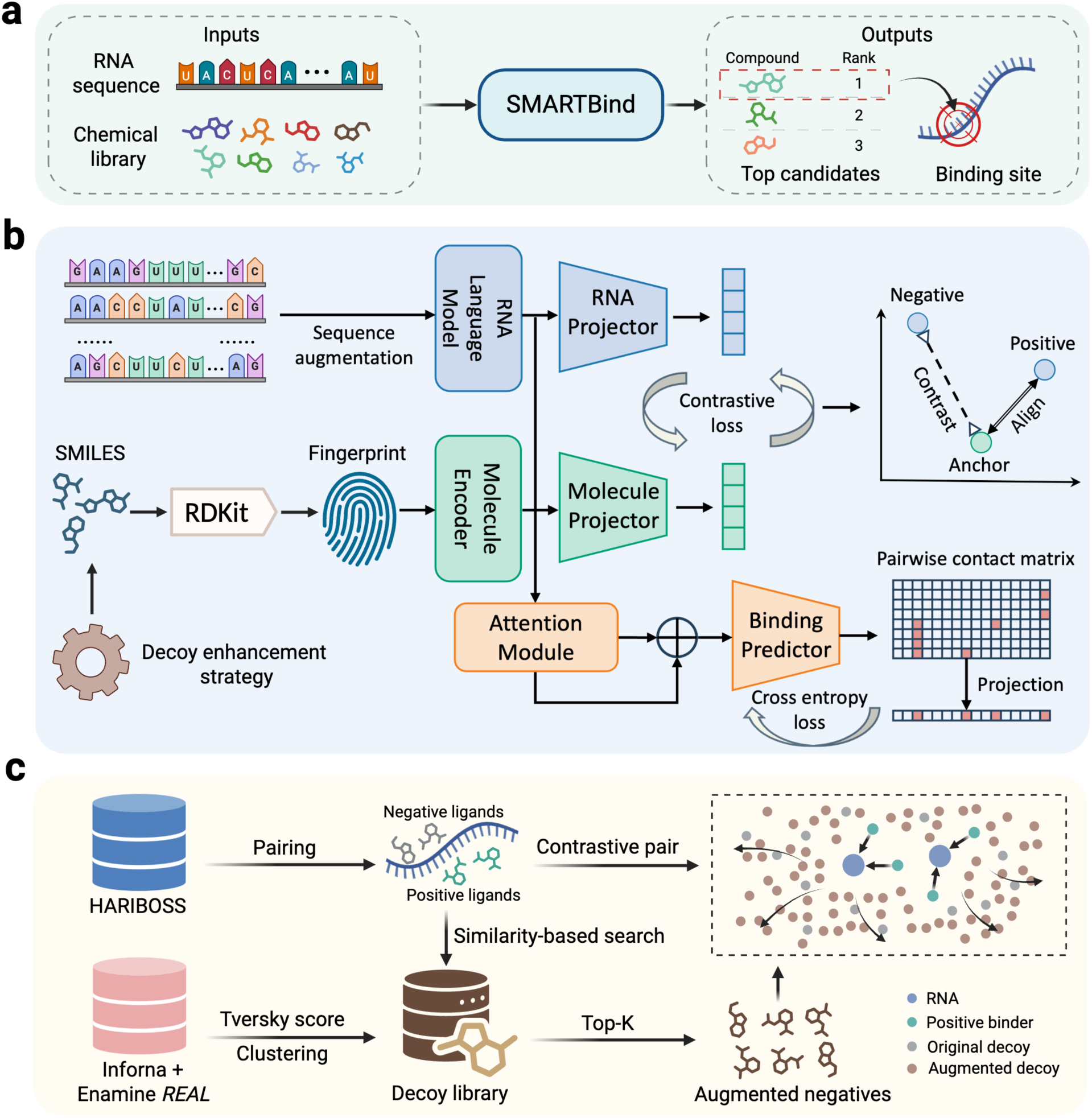
Overview of the SMARTBind framework for RNA targeting small molecule binder discovery. **(a)** Given a target RNA sequence as input, SMARTBind can accurately identify promising small molecule binders from a chemical library and predict their corresponding binding sites along the RNA sequence. **(b)** SMARTBind model architecture. RNA sequences are encoded by the pre-trained RNA language model to capture evolutionary and structural features, while ligands are represented as molecular fingerprints. Two projector modules align RNA and ligand embeddings in a shared latent space using a contrastive loss to distinguish binding from non-binding pairs augmented by the decoy enhancement strategy. A self-attention-based module predicts nucleotide-level binding sites from the encoder representations. **c,** Ligand-specific decoy enhancement strategy. A chemically diverse decoy library was constructed by screening bioactive molecules from the Inforna database against over 1 billion compounds in Enamine *REAL*, selecting candidates with Tversky similarity scores <0.4. A two-step K-means clustering reduced 2 million candidates to 92,626 structurally diverse compounds. During contrastive training, “challenging” decoys—those most similar to active binders—were dynamically selected for each RNA target to enhance the model’s discrimination ability and generalizability in virtual screening tasks.

Fig. 1b presents a schematic overall of the model architecture. The RNA sequence is processed using the pre-trained RNA language model RNA-FM^25^ to generate evolutionary and structurally informative feature representations. The ligand in SMARTBind is represented by FP2^26^, a 2D molecular fingerprint selected for its robust performance demonstrated in our benchmark and ablation studies (Supplementary Note 1). For the binding score prediction task, two projector modules are employed to transform the encoded representations of the RNA and small molecule into a unified latent space independently. In this space, the model is trained to align the embeddings of binding RNAs and small molecules while simultaneously contrasting non-binding molecules by optimizing the contrastive loss^27, 28^. The binding site prediction module is trained exclusively on RNA-ligand pairs, using representations encoded by the frozen base model pretrained for binding score prediction. The nucleotides involved in the interactions are identified by a self-attention-based predictor^29^.

To further improve SMARTBind’s robustness and generalization as a practical virtual screening application, a novel decoy enhancement strategy was developed that effectively expands and diversifies the model’s coverage of chemical space during training (Fig. 1c). A chemically diverse RNA-focused decoy library was first curated by screening all molecules from the Inforna^30^ database against a diverse set of drug-like compounds from Enamine *REAL*^31^ database (over 1.4 billion small molecules). Compounds with low Tversky similarity scores^32^ (<0.4) on molecular fingerprints were selected, followed by two-step clustering to identify 92,626 cluster centroids (∼15,000 small molecule per cluster), forming the final structurally diverse decoy library. The detailed construction process is provided in Methods and Supplementary Note 2. During the contrastive learning phase, compounds with the highest fingerprint similarity to the corresponding active binder were selected as “challenging” decoys for each specific RNA target (Fig. 1c). The average Tanimoto similarity^33^ calculated using FP2 fingerprint between decoys and their associated actives are 0.478 ± 0.101 (mean ± standard deviation [s.d.]), with the distribution shown in Supplementary Fig. 1. This strategy substantially enhances the model’s discrimination ability and generalizability in both benchmarking experiments and large-scale, real-world drug discovery tasks, *vide infra*.

### Benchmarking SMARTBind for RNA-ligand interaction prediction

SMARTBind was first evaluated on the RNAmigos1-curated dataset using its 10- fold cross-validation protocol^34^. All the RNA-ligand pairs in this dataset were collected from the Protein Data Bank (PDB)^35^. In the evaluation, true binders were embedded within a screening library, and their rank percentiles relative to the background decoy compounds were used as the primary performance metric. An ideal binder discovery model should achieve a rank percentile close to 1, indicating that the active binder is ranked near the top among all candidate compounds, while a random predictor will yield an average rank percentile of 0.5. Following the RNAmigos1 benchmark setting, two screening libraries were used: (1) the PDB Ligand set, derived from the test set by treating all ligands that do not bind the target RNA as decoys; and (2) the DecoyFinder set, generated from the ZINC^36^ database using DecoyFinder^37^ with default settings. For each true binder, DecoyFinder sampled 36 decoys that share similar physicochemical properties to the active ligand (see Methods), while maintinaing chemical dissimilarity (Tanimoto similarity ≤0.75). As shown in Fig. 2a, when evaluated on the PDB Ligand set, our model achieved an average rank percentile of 0.802, significantly outperforming the RNAmigos1 baseline (0.715). Similarly, on the DecoyFinder^37^ set, SMARTBind (0.821) consistently outperformed the baseline (0.735) by a substantial margin (Fig. 2a).

**Fig. 2.**
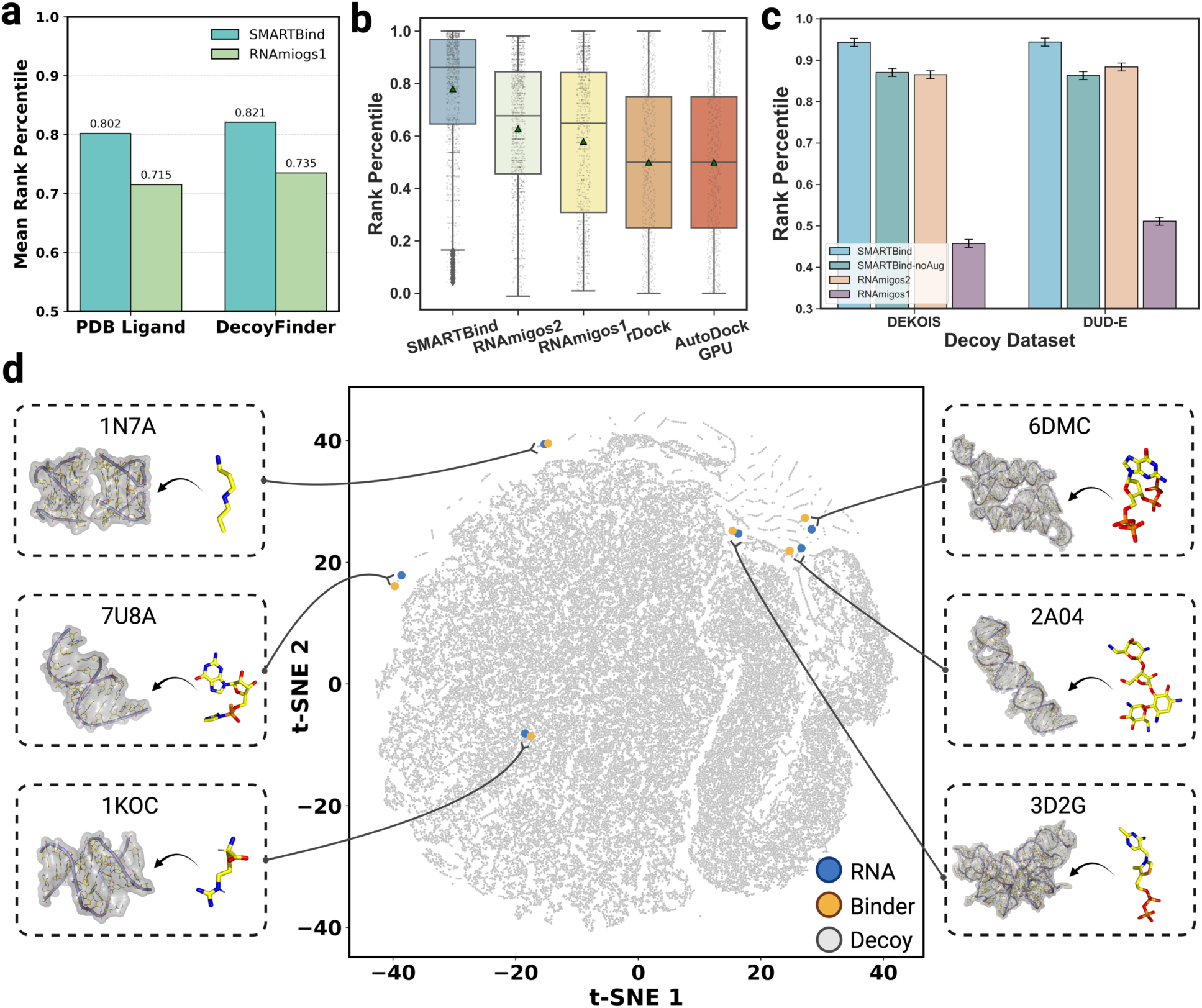
Performance comparison for RNA–ligand interaction prediction and analysis of SMARTBind’s latent space. **(a)** Performance comparison of SMARTBind and RNAmigos1 on the RNAmigos1-curated 10-fold random split dataset. Bar plots show mean rank percentiles for the PDB Ligand- and DecoyFinder-based screening libraries. **(b)** Performance comparison on the HARIBOSS 5-fold sequence-based split dataset using the PDB Ligand-based screening library. Box plots show the distribution of rank percentiles for SMARTBind, data-driven methods RNAmigos1 and RNAmigos2, and conventional docking methods rDock and AutoDock-GPU in a semi-blind docking approach. Dots represent individual positive RNA-ligand pairs, and green triangles indicate the mean rank percentile for each model. **(c)** Performance comparison on the HARIBOSS 10-fold random split dataset using two screening libraries generated by DeepCoy models trained on DEKOIS and DUD-E. Error bars represent the standard error of the mean across multiple folds. **(d)** t-SNE visualization of RNA and small moleucle co-embedding space. Grey dots represent decoy molecules, while blue and yellow dots denote RNA targets and SMARTBind-predicted binding ligands, respectively. Positive RNA–ligand pairs form distinct clusters against the molecular library background.

Our model was next applied to the HARIBOSS dataset^38^, using a more rigorous 5- fold cross-validation based on sequence similarity, ensuring that no RNA targets in the training and test sets share more than 30% sequence identity. Following the same evaluation procedure as above and using its associated PDB Ligand set as the screening library, we compared SMARTBind against data-driven approaches, including the RNAmigos series^34, 39^ and conventional docking methods including rDock^40^ and AutoDock-GPU^4^. Notably, all the comparison methods require either the detailed 3D structure of RNA or a 2.5D graph representation encoding canonical (Watson–Crick and wobble) base-pairing and non-canonical pairing interactions. In contrast, the structure-agnostic SMARTBind approach operates solely on the RNA’s sequence. Despite the absence of structural information, SMARTBind outperformed each baseline, achieving the highest average rank percentiles (0.779 ± 0.006 (mean ± standard error of the mean [s.e.m.]) compared to 0.627 ± 0.007 for the second-best model, RNAmigos2 (Fig. 2b), demonstrating a clear performance advantage.

To further assess the model’s screening capabilities, a larger screening library was created that was enriched with decoys generated by DeepCoy^41^, a deep learning-based model that designs physicochemical property-matched but structurally dissimilar molecules to the active binders. Two variants of the DeepCoy model, respectively trained on DEKOIS 2.0^42^ and DUD-E^43^, were used to generate 1,000 decoys per true binder in the HARIBOSS dataset. We compared SMARTBind with RNAmigos model series, as they demonstrated stronger performance over docking-based approaches in the preceding evaluations. As shown in Fig. 2c, SMARTBind achieved the highest rank percentiles on both the DEKOIS set (0.943 ± 0.006; mean ± s.e.m.) and the DUD-E set (0.944 ± 0.006), significantly outperforming the second-best model, RNAmigos2, which reached 0.865 ± 0.010 and 0.884 ± 0.010 on the two sets, respectively. We also evaluated SMARTBind- noAug, a non-augmented variant of SMARTBind that does not incorporate the decoy enhancement strategy during training. While it achieved performance comparable to RNAmigos2, with a rank percentile of 0.870 ± 0.008 on the DEKOIS and 0.863 ± 0.009 on DUD-E sets, it still notably underperformed relative to the standard SMARTBind model. The approximately 7% gain in rank percentile achieved by SMARTBind highlights the effectiveness of the decoy enhancement strategy in improving the model’s generalization ability for RNA-targeted ligand discovery. More comparison results of SMARTBind and SMARTBind-noAug can be found in Supplementary Note 3.

### SMARTBind captures an interaction-aware RNA-ligand co-embedding space

To elucidate the RNA-ligand co-embedding space established by SMARTBind for binding score prediction, t-SNE^44^ was utilized to visualize learned embeddings of RNAs and small molecules. The active pairs and decoys were randomly sampled from the test set, with RNA–ligand complexes sourced from the PDB and decoys from the curated decoy library. A clear clustering pattern was observed where true binders are located near their cognate RNA structures, such as SPM–1N7A, LXI–7U8A, ARG–1KOC, G4P–6DMC, NMY–2A04, and TPP–3D2G, while decoys exhibit a much more dispersed distribution across the embedding space (Fig. 2d). This demonstrates the model’s efficacy in capturing the RNA-small molecule interaction relationship, thereby rendering the co-embedding space interaction-aware. Furthermore, pair-wise cosine similarity scores were calculated to evaluate the distribution of embedding distances between RNAs and small molecules. As shown in Supplementary Fig. 2, the similarity scores are notably higher for true binding pairs compared to RNA-decoy pairs, with a one-sided Wilcoxon rank sum test (**p < 0.01, false discovery rate (FDR) corrected), confirming this distinction across both random and sequence-based split configurations. These findings provide valuable insights into the model’s predictive behavior.

Given that spatial relationships in the embedding space meaningfully reflect RNA– ligand binding interactions, SMARTBind enables prediction of their interactions based solely on the distances between RNA targets and small molecules mapped into this unified space. This mechanism significantly improves prediction efficiency by bypassing the time-consuming docking process, allowing rapid and scalable screening across large chemical libraries while maintaining high accuracy.

### SMARTBind enables accurate prediction of RNA-ligand binding sites

As a clustering pattern was observed for the true binders (Fig. 2D), SMARTBind’s ability to identify binding sites accurately given an RNA sequence and its corresponding small molecule binder was further evaluated. Binding nucleotides were defined as those with any heavy atom located within 10 Å of the ligand, and the area under the curve (AUC) was computed by comparing the model-predicted nucleotide-level binding probabilities with the binding site labels. As shown in Fig. 3a, the model delivers consistently accurate predictions of binding sites across both random split and sequence-based split settings, achieving average AUC scores of 0.674 ± 0.006 (mean ± s.e.m.) and 0.623 ± 0.006, respectively. The model’s performance across varying RNA chain lengths was further explored by categorizing the RNAs into three groups: short (<50 nt; on average 24 ± 11 nt, mean ± s.d.; n = 359), medium (50-200 nt; on average 86 ± 28 nt; n = 345), and long (>200 nt; on average 2187 ± 1298 nt; n = 542). The detailed length distribution is provided in Supplementary Fig. 3. Taking the random split as an example, SMARTBind achieved average AUC of 0.748 ± 0.013 (mean ± s.e.m.) and 0.737 ± 0.010 for short and medium-length RNAs, respectively (Fig. 3b). However, its performance decreased to 0.584 ± 0.007 for long RNA chains, such as ribosomal RNAs and riboswitches. This decline likely reflects the greater structural complexity and conformational heterogeneity of long RNAs, which often contain multiple potential binding pockets, in some cases intricate tertiary interactions, and dynamic conformational states^45^. Such characteristics remain challenging to capture fully at the sequence level, thereby increasing ambiguity in associating specific ligands with their true binding sites.

**Fig. 3.**
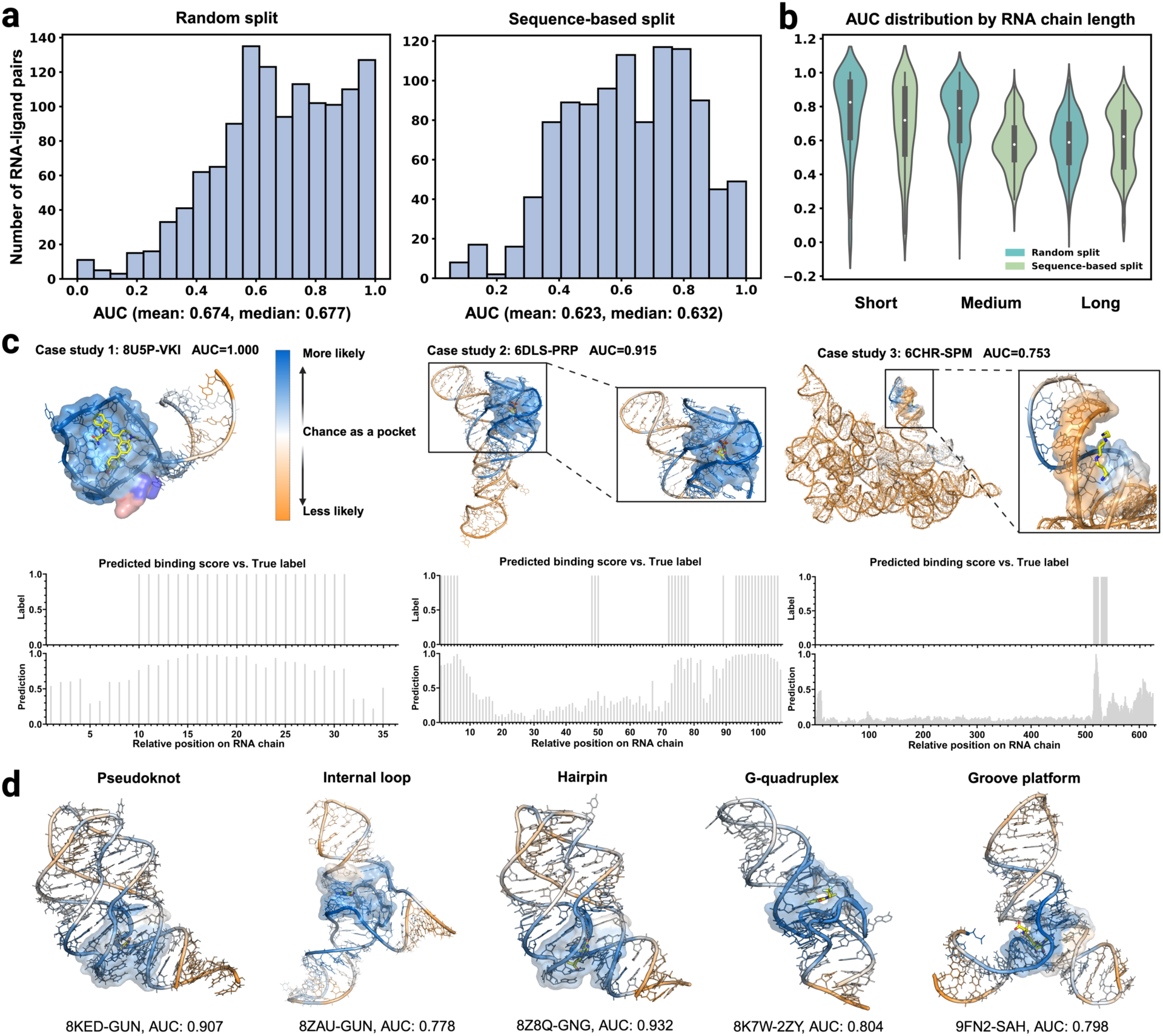
Assessment of binding site predictions with SMARTBind. **(a)** Distribution of AUC scores on the HAIRBOSS test set under two settings: 10-fold random split and 5-fold sequence-based split. **(b)** Distribution of AUC scores stratified by RNA sequence length: short (<50 nt), medium (50-200 nt), and long (>200 nt). **(c)** Binding site predictions of representative examples (PDBs: 8U5P, 6DLS, and 6CHR) from short, medium, and long RNA groups. The top panel illustrates structural visualization of predictions, with prediction confidence shown as a color gradient (blue for binding sites to orange for non-binding sites). Surfaces indicate reference binding sites. The bottom panel compares nucleotide-level predicted binding site probabilities with the ground truth labels. **(d)** Binding site predictions of representative examples from the temporally independent benchmark spanning diverse RNA structural classes: pseudoknot (8KED– GUN), hairpin (8Z8Q–GNG), internal loop (8ZAU–GUN), G-quadruplex (8K7W–2ZY), and groove platform (9FN2–SAH).

To further investigate model predictions, case studies on three representative RNA–ligand pairs from the test set under the sequence-based split settings were carried out, with one pair randomly selected from each of the three length-based categories: 8U5P (short), 6DLS (medium), and 6CHR (long). Fig. 3c illustrates the model’s normalized nucleotide-level binding site probability predictions, overlaid with the labels mapped onto the RNA sequences. For 8U5P and 6DLS, the predicted binding probabilities closely matched the experimentally determined binding nucleotides (AUC = 1.000 and 0.915, respectively), with high-probability regions delineating the ligand-binding pockets in 3D space. In contrast, the model did not conclusively identify the binding site for 6CHR (AUC = 0.753) from the 3D visualization. However, it still effectively narrows down potential regions, offering valuable guidance for elucidating the binding mechanism of the compound and supporting further experimental validation. These results demonstrate SMARTBind’s strong binding site prediction performance for short and medium RNAs, while revealing challenges for long RNAs.

In addition, performance on an external, temporally independent test set comprising 16 recent RNA–ligand complexes that were experimentally resolved and deposited in the PDB in 2025 (collected on 5 May 2025; Supplementary Table 1) was evaluated, after excluding any RNA targets present in the training set and removing non-interested small molecules (e.g. GTP, water molecules, and ions). This benchmark encompasses diverse binding motifs, including pseudoknots, hairpins, internal loops, G-quadruplexes, and groove platforms. As illustrated in Fig. 3d, SMARTBind accurately delineates ligand-binding sites across these varied architectures, achieving strong AUC values. Overall, the model achieved an average AUC of 0.749 ± 0.038 (mean ± s.e.m.; Supplementary Table 2), demonstrating robust generalization to recently resolved RNA– ligand complexes across diverse structural classes.

### SMARTBind prioritizes true binders for crucial RNA targets with confident binding site recognition

SMARTBind was next evaluated under more realistic ligand discovery scenarios, where active molecules are vastly outnumbered by decoys. To this end, a large-scale screening benchmark was constructed by randomly sampling one million small molecules from the Chemspace commercial library (over 7 million small molecules; See Methods), and spiking in a single experimentally validated RNA-binding ligand as the true positive (Fig. 4a). Models were assessed based on their ability to prioritize this true binder over a vast chemical background. For clarity of presentation, we report performance using the inverse rank percentile metric, in which compounds were sorted in descending order by predicted binding scores. In opposition to the above rank percentile metrics that use ascending order, lower inverse rank percentile values indicate higher prioritization of the true binder among the background compounds. We compared SMARTBind with existing data-driven methods including RNAmigos1^34^, RNAmigos2^39^, and the more recently published RNAsmol^46^, specifically RNAsmol_PDB and RNAsmol_ROBIN variants.

**Fig. 4.**
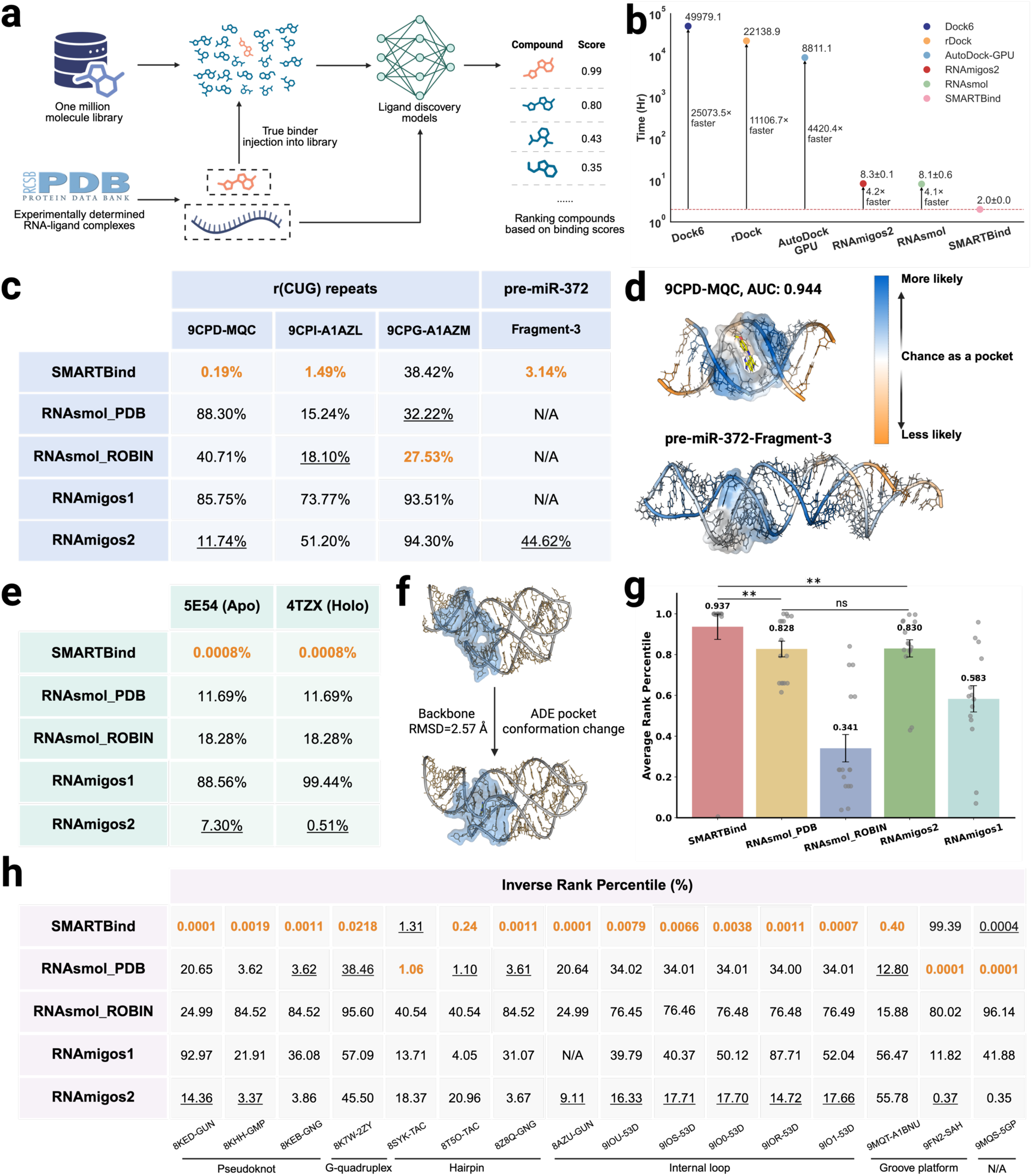
Evaluation of screening accuracy and efficiency in large-scale virtual screening case studies. **(a)** Schematic of the large-scale virtual screening benchmarking pipeline. A known RNA-ligand pair from the PDB was injected into a one-million compound library. Ligand discovery models were then tasked with screening the entire library and prioritizing the true binder. **(b)** Screening efficiency comparison on a 10-million-compound library. All methods, except AutoDock-GPU, were run on a single CPU core. Time is plotted on a logarithmic scale; error bars indicate standard deviation across five independent runs for SMARTBind, RNAsmol, and RNAmigos2. **(c)** Inverse rank percentile performance comparison on a single experimentally resolved RNA structure bound to three distinct active ligands (9CPD-MQC, 9CPI-A1AZL, 9CPG-A1AZM), as well as on pre-miR-372- Fragment-3, which lacks a resolved structure. Best results are highlighted in orange and shown in bold, the second-best results are underlined. **(d)** SMARTBind-predicted RNA– ligand binding sites for active binders. Prediction confidence is visualized as a color gradient from blue (binding site, high confidence) to orange (non-binding site, low confidence). **(e)** Inverse rank percentile performance comparison on an apo–holo RNA pair (PDBs: 5E54 and 4TZX) with substantial ligand-induced RNA conformational changes, evaluated against the holo-structure-bound ligand adenine riboswtich (ADE). Best results are highlighted in orange and shown in bold, the second-best results are underlined. **(f)** Structural visualization of the adenine-binding pocket in the apo and holo forms of the adenosine riboswitch. The pocket is defined as nucleotides within 10 Å of adenine in the holo structure, revealing conformational rearrangements with a global backbone RMSD of 2.57 Å. **(g)** Average rank percentile comparison for RNA–ligand complexes released in the PDB in 2025 (collected on 5 May 2025). Error bars represent standard error of the mean. Statistical significance was assessed using the Wilcoxon signed-rank test: **, *p* < 0.01 and not significant (ns). **(h)** Detailed inverse rank percentile performance for each target in (g). Best results are highlighted in orange and shown in bold, the second-best results are underlined.

The recently resolved RNA-small molecule structures^47^ (PDB IDs: 9CPD, 9CPI, and 9CPG, released on August 7, 2024) were used as the first set of case studies. These entries, which feature the same RNA target, r(CUG) repeat expansions implicated in Huntington’s disease-like 2 and myotonic dystrophy type 1, bound to three different small molecule ligands (MQC-9CPD, A1AZL-9CPI, A1AZM-9CPG). All were published after the training data utilized by SMARTBind, RNAmigos series and RNAsmol, thereby serving as unbiased, temporally separated test cases. As shown in Fig. 4c, SMARTBind ranked MQC and A1AZL at the top 0.19% and 1.49%, respectively, outperforming other methods; the second-best results were achieved by RNAmigos2 at 11.74% for MQC and RNAsmol_PDB at 15.24% for A1AZL. However, performance across all methods was suboptimal for A1AZM, where RNAsmol_ROBIN achieved the best ranking at 27.53%, compared to 38.42% for SMARTBind. A 10 Å cutoff from each ligand was applied to identify the corresponding binding nucleotides in each PDB entry, for evaluating the model’s ability to localize binding sites. SMARTBind achieved AUC scores of 0.944 for 9CPD (Fig. 4d, top), 0.786 for 9CPI (Supplementary Fig. 4a), and 0.847 for 9CPG (Supplementary Fig. 4b), demonstrating its ability to resolve RNA–ligand interactions at high resolution down to the binding nucleotide level.

In many early-stage drug discovery campaigns, RNA molecules of interest often lack experimentally determined 3D structures or are only available at low resolution. To illustrate SMARTBind’s practical applicability under such conditions, a case study on pre-miR-372, an RNA target lacking an experimentally determined 3D structure but with an in vitro validated binding fragment-3^48^ is presented. The RNA input to SMARTBind remains solely its primary sequence. In contrast, for the structure-based method RNAmigos2, we utilized FARFAR2 to generate high-confidence 3D structures and provided the extracted binding pocket, as determined by Inforna and in vitro binding assays^49^, as input. SMARTBind achieved the highest inverse rank percentile (3.14 %), while RNAmigos2 failed to identify the positive binder for pre-miR-372 effectively if given the predicted structure or pocket (Fig. 4c). This highlights the potential limitations of structure-based methods when relying on predicted RNA structures and pockets, in cases where experimentally resolved structures are unavailable. We additionally examined RNAmigos1 and RNAsmol; however, their respective implementations were unable to correctly parse the compound, resulting in “N/A” entries in the performance table. Moreover, SMARTBind’s predicted binding region encompassed the fragment-binding site of pre-miR-372 (an A-bulge previously identified^48^), as illustrated in the bottom panel of Fig. 4d, although the prediction covered a relatively broad area.

Even when crystal structures are available, binding-induced RNA conformational changes can also challenge current ligand discovery methods^50^. For example, riboswitches often exhibit substantial conformational changes between their “on” (unbound or apo) and “off” (ligand-bound or holo) states, and this conformational switching regulates downstream gene expression^51, 52^. We investigated model performance using an adenine riboswitch as a case study, where the apo (PDB: 5E54) and holo (PDB: 4TZX) forms exhibit a backbone RMSD of 2.57 Å (Fig. 4f). The corresponding RNA sequence was used as the input for SMARTBind, and it was absent from the model training data, ensuring an unbiased evaluation. For structure-based methods, the binding nucleotides for apo and holo were both defined based on the holo structure, using all nucleotides within 10 Å of the ligand as the input binding site. As shown in Fig. 4e, the second-best model, RNAmigos2 exhibited a notable performance decline (∼14-fold) on the apo structure (∼7.29%) compared to the holo structure (∼0.51%), highlighting the detrimental impact of binding-induced RNA conformational changes on the effectiveness of structure-based ligand discovery methods when only apo structures are available. In contrast, SMARTBind delivered consistent and robust top-ranking performance due to their structure-agnostic design, with SMARTBind remarkably ranking the ligand among the top eight out of over one million candidates. Although RNAsmol demonstrates robustness to conformational changes, both of its variants yielded significantly lower rankings (11.69% for RNAsmol_PDB and 18.28% for RNAsmol_ROBIN) compared to SMARTBind (0.0008%).

To further evaluate the model performance across more diverse RNA targets, we compared SMARTBind with other data-driven methods on the same temporally independent benchmark of 16 recently released RNA–ligand complexes (Supplementary Table 1). As shown in Fig. 4g, SMARTBind achieved highest average rank percentile across these unseen targets (0.937 ± 0.062, mean ± s.e.m.), substantially outperforming the other methods (RNAsmol_PDB: 0.828 ± 0.039, RNAsmol_ROBIN: 0.341 ± 0.067, RNAmigos2: 0.830±0.042, RNAmigos1: 0.583 ± 0.065). These improvements were statistically significant (p < 0.01) over the second-best models, RNAsmol_PDB and RNAmigos2. A refined analysis (Fig. 4h) revealed that SMARTBind consistently identified the true binder across a broad spectrum of RNA structural motifs. The benchmark encompasses pockets spanning diverse three-dimensional RNA motifs—including pseudoknots, G-quadruplexes, internal loops, groove platforms, and hairpins—as annotated by RNA 3D Hub^53^ (Supplementary Table 1).

Together, these results underscore the robustness and generalizability of SMARTBind in large-scale virtual screening applications. SMARTBind not only outperforms existing structure-based screening methods but also effectively addresses their key limitations, particularly in scenarios where RNA structural information is unavailable, low-confidence, or impacted by dynamic conformational changes.

### Ultra-efficiency on large-scale virtual screening

With the advent of ultra-large, make-on-demand compound libraries, efficiency has become a critical evaluation metric for VS approaches alongside accuracy. Thus, the screening speed of SMARTBind against other VS methods was measured. We first evaluated conventional docking-based approaches and observed that Dock 6 and rDock require 17.99 seconds and 7.97 seconds per compound on a single CPU core, while AutoDock-GPU takes 0.32 seconds per compound using a single GPU. It should be also noted that docking requires several preparatory steps before the actual docking can be performed. Scaling these estimates to a ten-million-compound library, Dock 6, rDock, and AutoDock-GPU would require approximately 5.71 years, 2.53 years and 1 year, respectively. In contrast, SMARTBind completed the same screening task in just 2.0 h on a single CPU core—demonstrating a dramatic improvement in efficiency and a substantial speed-up over docking-based methods (Fig. 4b). Under identical settings, the other data-driven approach, RNAmigos2 and RNAsmol, required 8.3 h and 8.1 h, respectively, to process the same dataset. Furthermore, since SMARTBind predicts RNA–ligand interactions based on the embedding distance between RNA targets and small molecules, its efficiency advantage can be further amplified when screening multiple RNA targets against the same compound library. Specifically, molecule embeddings can be precomputed once and stored in memory or a vector database, avoiding redundant calculations and costs and enabling efficient reuse across screening tasks.

Given the demonstrated ultra-efficient screening capability, SMARTBind is well-suited for exhaustive screening of billion-scale compound libraries within days, especially when leveraged with affordable parallel computing technologies. These results position the framework as a robust and scalable solution for RNA-targeted drug discovery, capable of triaging billions of compounds while maintaining accuracy and chemical diversity in selected hits, with potential to guide fragment-to-lead optimization in a data-driven, resource-efficient manner^54, 55^.

### Targeting pre-miR-21 with SMARTBind: discovery and validation of novel small molecule inhibitors

The maturation of oncogenic microRNAs (miRNAs) represents a promising therapeutic intervention point in cancer treatment, particularly for miR-21^56^, implicated as an oncomiR across diverse tumor types^57, 58^. As a post-transcriptional regulator, miR-21 represses numerous tumor suppressor genes, including Phosphatase and Tensin Homolog (PTEN) and Programmed Cell Death 4 (PDCD4), promoting cancer cell proliferation, survival, and invasion^59^. Elevated miR-21 expression is a hallmark of cancers such as glioblastoma, non-small cell lung cancer, and triple-negative breast cancer (TNBC)^60, 61^ (Fig. 5a). MiRNA biogenesis involves two sequential processing steps: conversion of the primary (pri-) miRNA to its precursor (pre-) form in the nucleus by Drosha. The pre-miRNA is then exported to the cytoplasm where it is further processed by Dicer to afford the mature miRNA^62^ (Fig. 5c). Mature miRNAs bind to the 3’ untranslated regions (UTRs) of mRNAs that share sequence complementarity and reduce gene expression by either inducing mRNA cleavage or translational repression.^63^ Blocking these biogenesis steps by binding to Drosha or Dicer processing sites offers a strategy to block miRNA maturation upstream and deactivate its oncogenic function^64^.

**Fig. 5.**
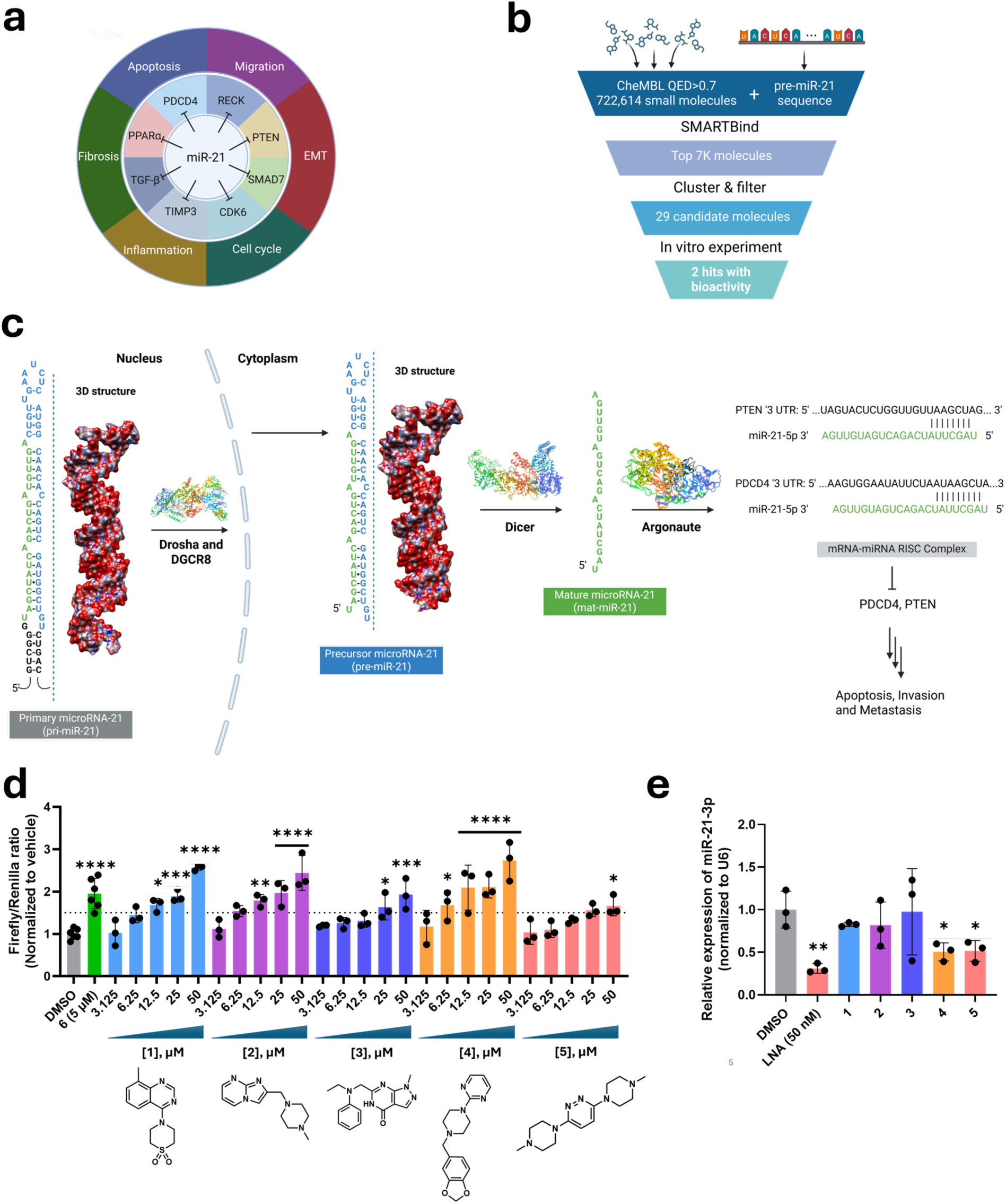
Integrated pipeline for the identification and functional validation of small-molecule inhibitors targeting miR-21. **(a)** Regulatory targets of miR-21 in various diseases and disorders, demonstrating the importance of this miRNA as diagnostic/prognostic marker and therapeutic target. **(b)** Virtual screening protocol for pre-miR-21. Screening was performed on a library of 722,614 drug-like small molecules, where drug-likeness was determined by a quantitative estimate of drug-likeness^65^ (QED) score ≥ 0.7. The top 1% of the library (n = 7,000) with the highest predicted binding scores was subjected to a two-step clustering process. The centroid of 100 resulting clusters was selected as candidate binders, and, of these, 29 were commercially available and procured for further analysis. Using only the RNA sequence of pre-miR-21, SMARTBind identified 29 structurally diverse candidate binders. **(c)** Schematics of pri-miR-21 biogenesis, including small molecule binding to inhibit Dicer processing and the interaction of the PDCD4 3’ UTR with miR-21. **(d)** Effect of compounds **1** – **5** on PTEN expression, as assessed using a luciferase reporter in MDA-MB-231 cells (n = 6 biological replicates for vehicle and the dovitinib derivative **6**; n = 3 biological replicates for SMARTBind compounds). Structures of all small molecules can be found in Supplementary Fig. 5a). **(e)** Effect of **1** – **5** (50 μM) on mature miR-21-3p abundance in MDA-MB-231 cells, as measured by RT-qPCR (n = 3 biological replicates for all treatment conditions). “LNA” indicates a miR-21-targeting LNA oligonucleotide (Supplementary Table 3). **, p < 0.01; ***, p < 0.001; and ****, p < 0.0001, as determined by a one-way ANOVA with multiple comparisons. Data are reported as the mean ± SD.

To evaluate the practical utility of SMARTBind in RNA-targeted ligand discovery, we conducted a virtual screening campaign against pre-miR-21, focusing on compounds predicted to bind to its Dicer processing site. Using only the pre-miR-21 sequence, SMARTBind screened a library of 722,614 drug-like small molecules ChEMBL library (2,496,335 compounds), where drug-likeness was determined by a quantitative estimate of drug-likeness^65^ (QED) score ≥ 0.7. The top 1% of the library (n = 7,000) with the highest predicted binding scores was subjected to a two-step clustering process. The first step of clustering was carried out by encoding a randomly sampled subset of the molecules (n = 1,000) using extended-connectivity fingerprints (ECFP4) and subjecting these fingerprints to multiple rounds of K-means clustering based on Tanimoto similarity. This afforded 155 cluster centers that were subsequently used to cluster the remaining 6,000 drug-like compounds. The centroid of 100 resulting clusters was selected as candidate binders, and, of these, 29 were commercially available and procured for further analysis (Supplementary Fig. 5a). The full screening pipeline is described in Fig. 5b.

Compounds prioritized by the model are predominantly fragment-like, suggesting an inherent bias toward smaller, less complex chemotypes. On one hand, this bias may be advantageous, as fragment-like molecules are often desirable starting points in fragment-based drug discovery due to their higher ligand efficiency, synthetic tractability, and potential for scaffold expansion.^66^ On the other hand, the tendency to favor fragments could also be limiting, as it may reduce chemical diversity and underrepresent larger, more drug-like scaffolds that might also yield biologically relevant interactions.

All 29 compounds were screened for reducing miR-21 levels using a luciferase reporter assay where the 3’ UTR of PTEN (direct downstream target of miR-21) is fused to firefly luciferase^58^. Thus, a small molecule that inhibits miR-21 biogenesis de-represses PTEN as measured by an increase in luciferase activity. Notably, the reporter is only responsive to miR-21 as the miR-17 binding site has been mutated. The PTEN luciferase reporter was co-transfected into MDA-MB-231 TNBC cells with a plasmid encoding Renilla luciferase, which was used as an internal control. The effect of the 29 compounds at two concentrations (5 µM and 50 µM) on PTEN expression was then measured using luciferase. Of the 29 candidates, ten compounds significantly increased firefly luciferase signal to levels comparable (>1.5 fold) to previously validated inhibitors of miR-21 biogenesis, Dovitinib^67^ and a derivative thereof (**6**)^68^, without affecting Renilla luciferase signal. Among these, five compounds (**1 – 5**) that increased firefly luciferase activity to while not affecting cell viability (Supplementary Fig. 5b-c) were selected for dose-response studies. [Cell viability assays indicated all compounds were tolerated at 5 µM, while eight reduced cell viability by >25% at the 50 µM dose (Supplementary Fig. 5b).]

While all five small molecules exhibited dose-dependent de-repression of PTEN, **1**, **2**, and **4** had similar profiles, increasing firefly luciferase signal by 2.13 ± 0.3-fold (p < 0.0001), 2.40 ± 0.23-fold (p < 0.0001), and 1.79 ± 0.22-fold (p < 0.0001), respectively, at the 50 µM dose (Fig. 5d). Some differentiation between compound activity was observed at the 5 µM dose where **1**, **2**, and **4** increased luciferase activity by 1.76 ± 0.15-fold (p < 0.0001), 2.28 ± 0.29-fold (p < 0.0001), and 1.74 ± 0.16 (p < 0.0001), respectively. In contrast, **3** and **5** were less potent, giving a 1.74 ± 0.58 (p < 0.0005) and a 1.71 ± 0.50-fold (p < 0.0005) signal increase at the 50 µM dose, respectively, and no observed activity at lower concentrations (Fig. 5d). To bolster confidence that the observed activity was due to an on-target effect, the five small molecules were tested in the same luciferase assay in HEK-293T cells. HEK-293T cells (C_t_ = ∼30) express much lower levels of miR-21 compared to MDA-MB-231 cells (C_t_ = ∼22; ∼256-fold higher expression)^59^ and thus were expected to be inactive. Indeed, no effect on PTEN expression (firefly luciferase activity) was observed for any of the five compounds (Supplementary Fig. 5d).

To evaluate whether the five SMARTBind-identified compounds de-repress PTEN expression by inhibiting miR-21 biogenesis, their effects on mature miR-21 levels upon treatment of MDA-MB-231 cells (50 µM, 48 h) was measured, as quantified by RT-qPCR quantification (Supplementary Table 3). Notably, **4** and **5** reduced mature miR-21 levels, by 49 ± 14% and 48 ± 14% (p < 0.05), respectively. In contrast, a very small (and statistically insignificant) effect was observed for **1** and **2,** while **3** was inactive. As **1** – **3** increased luciferase activity in the reporter assay (Fig. 5d), these results suggest that they may have a mixed mode of action.

The reduction of mature miR-21 abundance by **4** and **5** was further characterized by measuring pre- and mature miR-21 levels in dose response after a 48 h-treatment period. For **4**, reduction of mature miR-21 abundance was only observed at the 50 µM dose, by 55 ± 5% (p < 0.005), in line with the single dose screen of the five molecules (Fig. 6a and Supplementary Fig. 5c). In contrast, **5** showed dose dependent reduction of mature miR-21 levels, by 43 ± 5% (p < 0.01) at the 50 µM dose and 30 ± 8% (p < 0.05) at the 20 µM dose. A dose-dependent increase in pre-miR-21 levels was observed for both **4** and **5**, where they were increased by 70 ± 8% (p < 0.01) and 56 ± 13% (p < 0.05), respectively at the 50 µM dose (Fig. 6b). These results are consistent with the hypothesis that **4** and **5** inhibit Dicer processing of pre-miR-21 and thereby de-repress PTEN in the luciferase reporter assay (Fig. 5d).

**Fig. 6.**
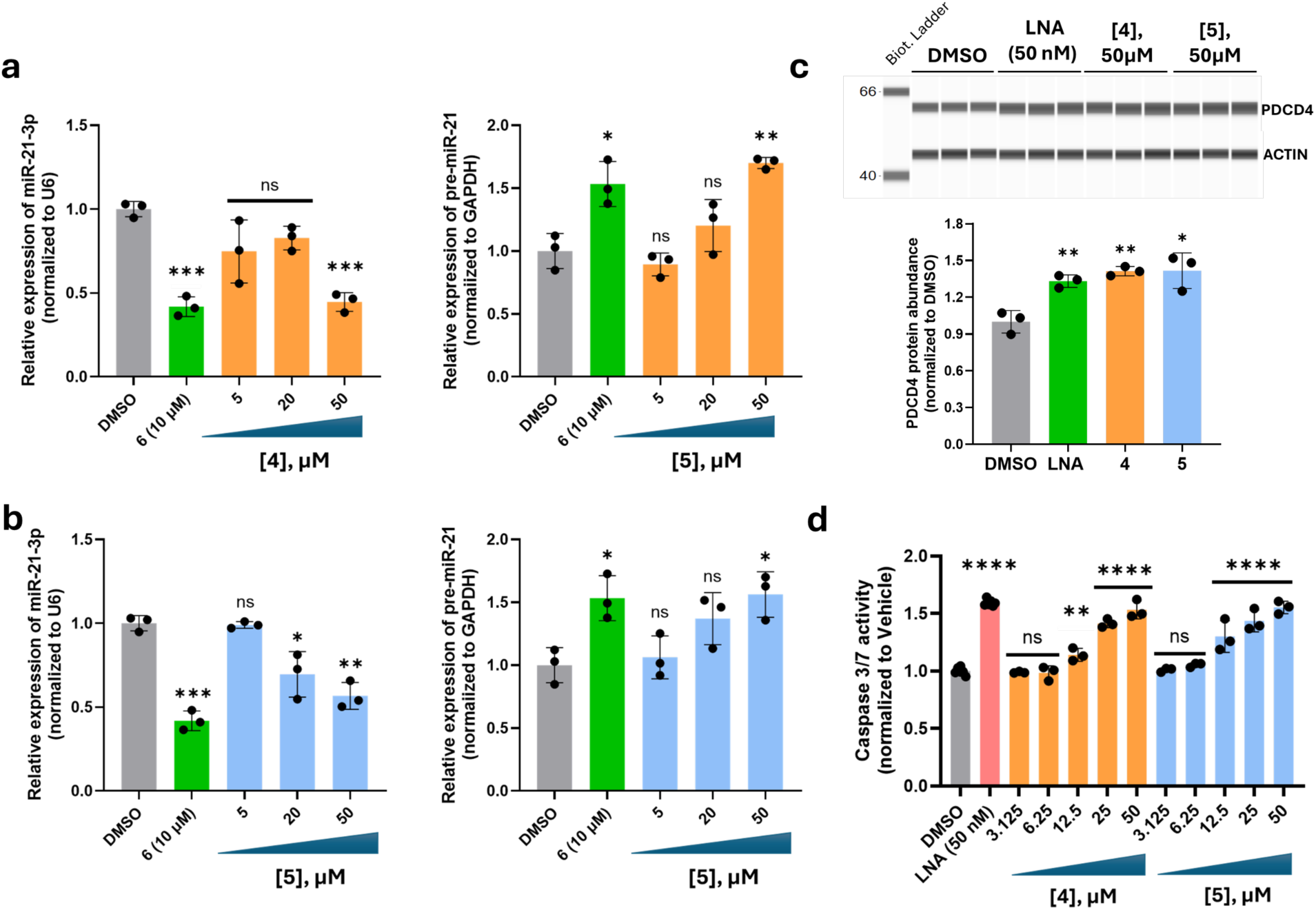
Compounds 4 and 5 inhibit miR-21 biogenesis in MDA-MB-231 cells, de-repress endogenous Programmed Cell Death 4 (PDCD4) expression, and promote apoptosis. **(a)** Effect of **4** on mature miR-21-3p and pre-miR_21 levels upon treatment of MDA-MB- 231 cells, as assessed by RT-qPCR (n = 3 biological replicates for all treated samples). **(b)** Effect of **5** on mature miR-21-3p and pre-miR-21 levels upon treatment of MDA-MB-231 cells, as assessed by RT-qPCR (n = 3 biological replicates for all treated samples). For panels a and b, **6** is a previously validated inhibitor for miR-21 biogenesis and is an analog of dovitinib^68^. **(c)** Representative Simple Western and quantification of PDCD4 abundance in MDA-MB-231 cells treated with **4** or **5** (50 μM; n = 3 biological replicates). **(d)** Effect of **4** and **5** on Caspase 3/7 activity upon treatment of MDA-MB-231 cells, as assessed by Caspase-Glo 3/7 Assay (n = 3 biological replicates). “LNA” indicates a miR- 21-targeting LNA oligonucleotide. **, p < 0.01; ***, p < 0.001; ****, p < 0.0001, as determined by a one-way ANOVA with multiple comparisons. Data are reported as the mean ± SD.

The results from RT-qPCR and luciferase reporter assays suggest that small molecules **4** and **5** inhibit miR-21 biogenesis by binding to the Dicer processing site while results for the other small molecules suggest a mixed mode of action. To gain insight into these observations, the binding of each small molecule to a model of the miR-21 Dicer site (Supplementary Fig. 6a) was investigated by using nuclear magnetic resonance (NMR)- based methods. In particular, Carr–Purcell–Meiboom–Gill (CPMG)^69^ relaxation dispersion experiments were employed that assess changes in spectral intensity and relaxation behavior upon ligand addition (indicative of binding). In each case, 5 μM of the Dicer processing site was added to 300 μM of small molecule. As shown in Supplementary Fig. 6, binding was observed for all five small molecules, however changes in the CPMG spectra for **1** – **3** were quite modest, consistent with very small changes observed in mature miR-21 abundance upon compound treatment as measured by RT-qPCR. Significant changes were observed in the CMPG spectra upon addition of the miR-21 Dicer site for **4** and **5**. For the former, the aromatic protons between 6.8 to 7.0 ppm corresponding to the protons of the benzene ring of the benzodioxazole moiety and the pyrimidine ring were exchange broadened (decreased peak intensity). For **5**, addition of the Dicer site reduced the intensity of the peak at 7.26 ppm corresponding to the protons of the pyridazine ring and shifted the peak upfield by 0.04 ppm. Importantly, these changes were not observed upon addition of a fully paired control RNA (Supplementary Fig. 6d), indicating specificity for the miR-21 Dicer site.

The changes observed in CPMG spectra suggested that the dissociation constant (K_d_) for **4** and **5** might be measurable. To do so, chemical shift perturbations (CSPs) in the proton spectra of the ligand were measured as a function of RNA concentration^70^. For compound **4**, quantifying the aforementioned CSPs as function of RNA concentration afforded a K_d_ of 83 ± 14 µM. Likewise, fitting the changes in chemical shift of protons in the pyridazine ring of **5** afforded a K_d_ 84 ± 16 µM. As **4** and **5** are fragment-like, these dissociation constants in the micromolar range are expected.

As RT-qPCR studies showed inhibition of miR-21 biogenesis by **4** and **5**, their effects on the endogenous levels of PDCD4, a downstream target of miR-21,was assessed by Western blotting. PDCD4 suppresses apoptosis by inhibiting the translation of procaspase-3 mRNA^71^. Repression of PDCD4 expression by miR-21 promotes cancer cell proliferation and reduces apoptosis in many cancers^71^. In MDA-MB-231 cells, **4** and **5** (50 µM) de-repressed endogenous PDCD4 protein expression by 41 ± 6% (p < 0.01) and 42 ± 10% (p < 0.05), respectively, as measured by simple Western (Fig. 6c). Moreover, this de-repression of PDCD4 is sufficient to re-activate apoptosis, where both **4** and **5** dose-dependently increased Caspase-3/7 activity, reaching a maximum increase of 53 ± 4% (p < 0.0001) and 55 ± 3% (p < 0.0001), respectively, at the 50 µM dose.

Taken together, these results establish **4** and **5** as RNA-binding fragments that inhibit miR-21 biogenesis, de-repress tumor suppressors, and promote apoptosis in TNBC cells. While their binding affinities are modest (Supplementary Fig. 6) — a characteristic feature of small molecular fragments — both compounds demonstrate clear functional effects in cell-based assays, including the reduction of mature miR-21 levels, increase of pre-miR-21 abundance, de-repression of PTEN and PDCD4, and activation of Caspase-3/7. These data suggest a specific mechanism of action consistent with inhibition of Dicer processing.

Importantly, **4** and **5** possess physicochemical properties consistent with fragment-like molecules, including low molecular weight and minimal structural complexity. As such, they represent ideal starting points for structure-guided optimization. Their chemical tractability allows for rapid analog generation to improve potency, selectivity, and pharmacokinetic profiles. Future medicinal chemistry efforts can leverage these scaffolds to design more potent and lead-like ligands that retain binding to the RNA target while maintaining drug-like properties. These efforts would focus on improving the activity of compounds and a more thorough assessment of the selectivity, all key requirements to turn these proof-of-concept compounds into lead medicines targeting disease-relevant pre-miR-21.

As an application of SMARTBind to a real-world RNA-targeted small molecule discovery project, this study not only validates the platform’s predictive utility but also highlights the feasibility of using fragment-based virtual screening to identify chemically tractable leads for RNA-targeting therapeutics. These findings support a broader strategy in which early hits — even those with relatively low binding affinities — serve as foundational cores for lead development campaigns aimed at RNA targets once deemed undruggable.

## DISCUSSION

RNA-targeted small molecules offer a promising therapeutic strategy for diseases that remain inaccessible to protein-targeting approaches, including genetic disorders, cancers, and viral infections. Virtual screening has emerged as a key component of early-stage drug discovery, allowing for the efficient and cost-effective exploration of vast chemical libraries^72, 73^. While virtual screening has been extensively applied to protein targets, its adaptation to RNA lags significantly behind. Both classical docking-based and newer data-driven, docking-free methods often show limited performance in the RNA context^74, 75^. This is largely due to the intrinsic challenges of RNA as a drug target, including its highly dynamic, solvent-exposed binding surfaces, noncanonical base pairing, shallow grooves, and flexible loop architectures—features that are not easily accommodated by traditional rigid scoring functions^76, 77^.

To overcome these limitations, we developed SMARTBind, a structure-agnostic, sequence-based RNA-targeted ligand discovery framework that predicts small molecule binding and localizes binding sites using only primary RNA sequence. Benchmarking across a diverse range of RNA targets and virtual screening tasks demonstrates that SMARTBind consistently outperforms state-of-the-art structure-based approaches. By leveraging foundation model-derived RNA sequence embeddings, RNA-ligand contrastive learning, and a ligand-specific decoy enhancement strategy, SMARTBind improves the hit discovery performance, all while eliminating the need for experimentally determined or predicted RNA structures (Figs. 2 and 4). Moreover, it substantially reduces computational time and resource demands, enabling ultra-large-scale virtual screening of millions to billions of compounds within hours on modest hardware—an efficiency orders of magnitude greater than that of traditional docking pipelines.

To validate SMARTBind’s utility in a real-world application, it was applied to pre-miR-21, a well-characterized oncogenic RNA involved in post-transcriptional regulation of tumor suppressors in triple-negative breast cancer^69, 78^. From an initial screen of drug-like small molecules, SMARTBind prioritized several candidates, five of which demonstrated dose-dependent de-repression of PTEN in a luciferase reporter assay (Fig. 4). Two compounds were found to inhibit miR-21 biogenesis, de-repress endogenous PDCD4, and promote apoptosis in TNBC cells (Fig. 5). Although the top-performing compound is well-tolerated and biologically active, it possesses physicochemical features characteristic of low molecular weight fragments. Accordingly, further lead optimization efforts are underway, including structure-guided analog design, fragment expansion, and conjugation to effector-recruiting modules, with the goal of enhancing potency and improving drug-like properties. These findings confirm SMARTBind’s ability to identify bioactive RNA-binding compounds directly from sequence input and highlight its practical relevance to fragment-based screening and medicinal chemistry workflows.

Compared to existing virtual screening methods, SMARTBind provides several critical advantages. First, traditional docking pipelines are complex, multi-step procedures involving structure prediction, binding site identification, docking, and scoring. Each of these steps introduces potential inaccuracies that can accumulate and reduce overall screening success, particularly for RNA where methodological development is less mature than for proteins^79^. SMARTBind replaces this fragmented approach with a single, end-to-end predictive model, reducing the opportunity for error propagation. Second, many data-driven docking-free approaches still rely on RNA structural input, making them inherently vulnerable to inaccuracies due to the dynamic, charged nature of RNA and the limited availability of high-quality structural data. In contrast, SMARTBind eliminates the need for RNA structure altogether by operating directly on RNA sequence, broadening the scope of targetable RNAs and facilitating learning from large, unlabeled sequence datasets. Third, as compound libraries continue to grow at an exponential rate—e.g., Enamine *REAL* (10.1 billion compounds) and ZINC- 22^23^ (54.9 billion compounds)—computational cost is becoming a key bottleneck. Even with access to parallel cloud computing resources, traditional docking becomes impractical at this scale^43^. SMARTBind addresses this challenge by using fast, embedding-based distance calculations between RNAs and small molecules, avoiding expensive ligand conformational sampling and enabling brute-force screening of ultra-large libraries.

Despite its strengths, SMARTBind has limitations. It still faces challenges in achieving precise binding site localization, particularly for long-length RNAs, highlighting the need for enhanced strategies such as hybrid approaches that integrate data-driven learning with physicochemical descriptors and RNA secondary or tertiary structural annotations. Furthermore, although high-confidence decoys were computationally generated to train the contrastive learning framework, incorporating experimentally validated inactive ligands when available would likely improve model accuracy. Lastly, the field would benefit from standardized, unbiased RNA–ligand benchmark datasets to allow rigorous cross-comparisons of scoring functions and virtual screening methods.

In conclusion, the ability to accurately identify small molecule binders to RNA is fundamental to advancing our understanding of RNA biology and to realizing the therapeutic potential of RNA-targeted small molecules. SMARTBind provides a robust, scalable, and structure-independent framework for RNA-specific ligand discovery, with demonstrated utility in both benchmark and real-world scenarios. Its unique combination of accuracy, speed, and generalizability positions it as a powerful tool for the discovery of next-generation RNA-targeted therapeutics.

## ACKNOWLEDGEMENT

This work was supported in part by the University of Florida (UF Startup Fund, UF Health Cancer Center Pilot Grant # UFS-2023-08, and UF Research AI Award to Y. L.), National Institutes of Health (R01 CA249180 to M.D.D.), the Florida Department of Health (Florida Cancer Innovation Fund grant #MOABO to J. L. C.), and the Muscular Dystrophy Association (Development grant ID #963835 to A. T.). Figures were created in Biorender https://BioRender.com.

## METHODS

### Dataset preparation

#### Training dataset preprocessing

The HARIBOSS database^38^ was used as one of the primary benchmark datasets for training and evaluating the performance of RNA-ligand interaction prediction models. All entries in the HARIBOSS were curated from the Protein Data Bank (PDB) and consist of experimentally validated crystal structures of RNA–drug-like compound complexes. As of this study, the database includes 862 RNA-small molecule complexes, 1,471 annotated binding pockets, and 311 unqiue ligands. The drug-like ligand natrually serves as the active binder for the correponding RNA target. For the binding site annotation, RNA nucleotides were defined as interacting if any of their atoms were located within 10 Å from the ligand atoms. We also identified several entries where multiple identical ligands bind to different regions on the same RNA chain. To annotate all potential binding sites for each unqiue RNA-ligand pair, we merged the binding sites corresponding to the same RNA sequences and ligands. The resulting dataset of 1,246 unique RNA–ligand samples was subsequently splitted by different stragegies for comprehensive benchmark purposes. For a fair comparison with the RNAmigos1 baseline, we also utilzed its curated dataset^34^, which similarily comprises RNA-ligand complex 3D structures from the PDB but was collected earlier and subjected to different filtering criteria than HARIBOSS. For example, binding sites with fewer than five RNA nucleotides were removed. This resulted in a dataset consisting of 773 binding sites associated with 270 unique ligands.

To comprehensively benchmark the methods, we adopted two distinct data splitting. First, for RNAmigos1 dataset, we followed its original 10-fold random split protocol to ensure consistency in comparison. The same protocal was also applied for the HARIBOSS dataset for parallel benchmarking. However, random splitting may introduce information leakage, where similar RNA seqeunces appear in both the training and test sets, potentially leading to overly optimistic performance estimates for machine learning-based methods. To mitigate this issue, we employed an RNA sequence identity-based method, as an alternative spliting protocol on the HARIBOSS dataset. Specifically, RNA chains were clustered at 30% sequence identity using MMseqs2^80^, and the resulting non-redundant clusters were then partitioned into five folds for cross-validation (Supplementary Fig. 7). The largest cluster encompassed nearly one-fifth of the dataset, motivating the use of a 5-fold sequence-based split for cross-validation to enable a more rigorous assessment of model generalizability.

#### Evaluation screening library curation

To evaluate the effectiveness of ligand discovery methods, we curated diverse and representative screening libraries for each RNA target in the benchmark test set. Each library comprised one positive sample (the true binder) together with a set of decoys generated through different strategies.

For the RNAmigos1 test set, we followed the authors’ protocol^34^ to generate decoy sets using two stratgies: (1) the PDB Ligand set in which all other ligands in the test set— excluding the known active binder—served as decoys for each RNA target; and (2) the DecoyFinder^37^-generated set, where 36 decoys per compound were sampled from the ZINC database^36^. Compounds in DecoyFinder-generated set were selected to share similar generic physical properties with the active binder—including molecular weight, number of rotatable bonds, total hydrogen bond donors, total hydrogen bond acceptors, and the octanol–water partition coefficient (logP)—while remaining chemically dissimilar (Tanimoto similarity ≤ 0.75), thereby potentially perturbing binding potential.

When benchmarking on the HARIBOSS test set, in addition to adopting the same protocol to curate the PDB Ligand set, we also employed DeepCoy, a deep learning-based method, to generate property-matched decoys that closely resemble the physicochemical profiles of active molecules while being structurally dissimilar^41^. Unlike database-dependent methods such as DecoyFinder, which may fail to identify sufficiently diverse decoys due to limitations of the search database, DeepCoy generates compounds de novo, enabling scalable expansion of decoy sets. Two variants of the DeepCoy model—trained on DEKOIS 2.0^42^ and DUD-E^43^, respectively, and released by the authors—were used directly to generate 1,000 decoys for each true binder in the HARIBOSS dataset. The average Tanimoto similarity calculated using FP2 fingerprint between decoys and their associated actives are 0.152 ± 0.055 (mean ± standard deviation [s.d.]) for DEKOIS 2.0- trained model and 0.191 ± 0.082 for DUD-E-trained model, with the distribution shown in Supplementary Fig. 8. The DUD-E-trained model was optimized by the authors to match 27 physicochemical descriptors, while the DEKOIS 2.0 version focused on eight molecular features, including molecular weight, LogP, number of hydrogen bond acceptors, number of hydrogen bond donors, number of rotatable bonds, number of aromatic rings, positive charge, and negative charge. A detailed description of these features is available in ref.^41^.

To benchmark the accuracy and efficiency in real-world case studies, we curated large-scale screening libraries from the Chemspace commercial database (7,456,971 entries)—a comprehensive collection of synthetically accessible molecules widely utilized in drug discovery. For accuracy evaluation, one million small molecules were randomly sampled from the Chemspace database to construct the chemical background set, together with one experimentally validated true binder. For the efficiency benchmark, the screening library comprised ten million compounds randomly drawn from the Chemspace library.

To construct a temporally independent benchmark, we curated RNA–ligand pairs from the PDB (downloaded on 5 May 2025) and retained only those released after 31 December 2024. We further excluded pairs whose RNA sequences appeared in the SMARTBind training set or whose ligands were GTP, water molecules, or ions, resulting in 16 PDB entries for evaluation. The pocket 3D motifs were annotated by using the RNA 3D Hub^53^. For sequence-based methods, including SMARTBind and RNAsmol, inference was performed directly from the full RNA sequence and ligand SMILE. For the structure-based RNAmigos series, RNA pockets within 10 Å of the bound ligand and the ligand SMILES were provided as input for inference.

#### SMARTBind framework

To formulate the problem, we represent the RNA sequence of interest as *r*, and a set of *n* small molecules as *M* = {*m*_1_, *m*_2_, …, *m*_*n*_}. Our binding scoring prediction task is to develop a scoring function *s*(·, ·) to predict the likelihood of binding to the RNA target *r* given small molecule *m*_*i*_. This scoring function *s*(·, ·) can then be used to guide the ligand discovery process, with the objective of identifying the top *k* molecules with the highest likelihood of binding to the target RNA. The binding site prediction module is designed to localize the most probable binding regions on the RNA target *r* for a given active binder *m*_*i*_, by predicting a nucleotide-level probability distribution over the RNA sequence, thereby offering additional insights into the binding mechanism.

#### Input processing

##### RNA featurization

Multiple RNA foundation models have been proposed to improve our understanding of RNA structures and functions. In this study, we utilized the RNA-FM^25^, a robust foundation model based on the BERT architecture^81^, as our RNA encoder. RNA-FM was pre-trained on 23 million unlabelled non-coding RNA (ncRNA) sequences by reconstructing the masked nucleotides from the sequence context alone^25, 81^. Taking the nucleotide sequence *r* of length *L* as input, RNA-FM generates an embedding matrix of dimensions *L* × 640, which is expected to encapsulate rich information from the ncRNA universe. To obtain a global and consolidated representation of the entire sequence, the embedding matrix was aggregated by computing the mean of each nucleotide embedding across the sequence, resulting in a single 640-dimensional vector. Average pooling is chosen as the aggregation mechanism because it is widely used, unbiased, and conservative, while also offering efficiency and simplicity^82, 83^. Subsequently, a non-linear projection model, consisting of a multi-layer perceptron (MLP) with LeakyReLU activation function^84^, is used to project this vector into the RNA-ligand co-embedding space *R*^ℎ^, where ℎ = 256, and the RNA embedding is denoted as *x*^*r*^. All hyperparameters reported in this and subsequent sections were determined empirically on the basis of benchmarking experiments.

##### Small molecule featurization

We explored multiple strategies for featurizing the small molecules into fixed-dimension embeddings *m* ∈ *R*^*dm*^, including pre-trained molecular foundation models such as MolCLR^85^, as well as widely used molecular fingerprints: FP2, Morgan, Topological Torsion (TT), Atom Pair (AP) and Extended 3- Dimensional Fingerprint (E3FP). Each featurization method has its unique advantages and disadvantages. We benchmarked their contributions to the SMARTBind-noAug version using HARIBOSS dataset with the 5-fold sequence-based split. Detailed information can be found in the Supplementary Note 1. Ultimately, we select the FP2 fingerprint, which encodes small molecule inputs as *d*_*m*_ = 1024-dimensional binary bit vectors, due to its consistently robust and superior performance in our benchmarking compared to other molecular fingerprints and pre-trained molecular foundation models. These features are then mapped to the RNA-ligand co-embedding space as a vector of *x*^*m*^ ∈ *R*^ℎ^ using another non-linear projection model.

##### Data augmentation

Data augmentation is crucial in prevalent unsupervised and supervised contrastive learning frameworks, effectively boosting model resilience by producing various augmented perspectives that preserve the semantic consistency of the original data^28, 86^. It has also proved to be an effective solution to data scarcity in low-resource scenarios. However, general augmentation strategies used for modelling biomolecular sequences, such as, random cropping, mutation, and masking, are are not always directly applicable to RNA-ligand interaction data. This is primarily because the key regions responsible for binding may vary in length and location across different RNA sequences. Applying such transformations indiscriminately can disrupt critical binding patterns, potentially breaking true interaction relationship and introducing noise into the positive training data. To alleviate such side effects, our data augmentation approach focuses on non-interacting regions, defined as nucleotides located at least 10 Å away from any atom of the ligand. During the training, every sequence will have 25% prossibility to be randomly mutated at approximatedly 15% of its non-contact nucleotides. This strategy introduces data variability while likely preserving the RNA’s structural integrity and functional relevance, which are essential for accurate binding interaction prediction. To enable efficient batch training with variable-length RNA sequences, long sequences were randomly cropped to 512 nucleotides with the constraint that the retained segment included at least one nucleotide in contact with the active binder. Sequences shorter than 512 were zero-padded.

#### Contrastive learning for RNA-ligand scoring function development

Given the RNA target embedding *x*^*r*^ ∈ *R*^ℎ^ and small molecule embedding *x*^*m*^ ∈ *R*^ℎ^, we first calculate the distance measurement 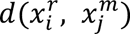 between each RNA-ligand pair 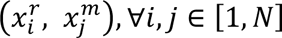. This distance measurement is inversely correlated with our defined binding scoring function 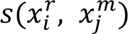, which is computed as the cosine similarity between their embedding vectors, followed by a sigmoid function *σ*:

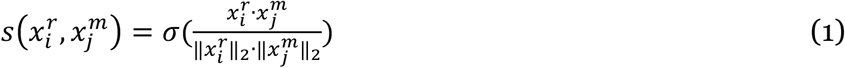

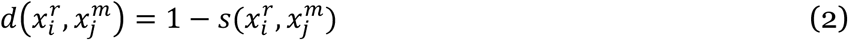

Contrastive learning is employed to train the scoring function network, aiming to minimize the distance between the positive RNA-ligand pairs while maximizing the distance between the negative pairs. Positive pairs naturally arise from the known RNA-ligand interaction data. However, we lack a well-established and experimentally validated dataset containing decoy samples for these RNA targets. Therefore, we construct our negative pairs inspired by the negative sampling strategy utilized in previous studies^27, 87^. Specifically, given the known RNA-ligand interaction data 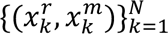, we first extract two lists consisting of RNAs 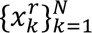 and small molecules 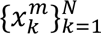, respectively. Then, we pair-wise combine an RNA and a small molecule from the two lists, resulting in *N*^2^ pairs 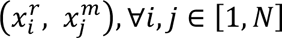. The positive pairs are the samples where *i* = *j*, and negative pairs are those where *i* ≠ *j* and no binding relationship is reported. This negative pair construction also aligns with the principle of decoy set construction used in baseline approaches like RNAmigos1^34^. Both methods are based on a general assumption that if a small molecule is an active binder to a specific RNA, it is unlikely to bind to other distinct RNAs, due to selective binding driven by specific RNA motifs, unique ligand-binding pockets, and higher-order structural features of both the RNA and the ligand. However, we argue that establishing an RNA-ligand dataset with experimentally verified binders and decoy samples is still crucial and urgent for the development of machine learning and deep learning methods.

The established positive and negative pairing data enable us to set up the contrastive learning objective with the required anchor, positive and negative training points. Based on different anchor types, we define two symmetric losses: RNA-anchor loss *L*_*rna*_ and ligand-anchor loss *L*_*li*g_. In the RNA-anchor loss, the anchor and positive instances are represented by an RNA target 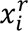 and its active binder 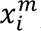, respectively. The negative samples are 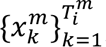, randomly selected from the negative pairs data 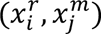, where *i* ≠ *j* and 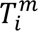 represents the number of sampled negative small molecules for 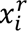. The triplet margin-distance loss is employed to shorten the distances between an anchor and its positive point while increasing the distances between the anchor and its negative examples, as follows:

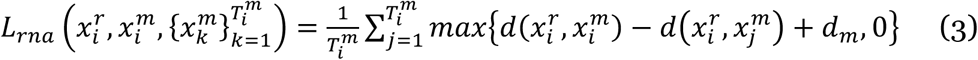

where *d*_*m*_ is a nonnegative margin parameter imposing the distance between the positive and negative samples to the anchor point to be larger than *d*_*m*_. This loss term describes the likelihood of a pool of small molecules binding to a given RNA target 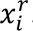.

In contrast, the ligand-anchor loss describes the likelihood of a pool of RNAs binding to a given small molecule, with 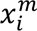 as the anchor. The positive and negative samples are its binding RNA target 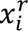 and any sampled non-binding ones 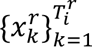, where 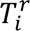 represents the number of negative RNA points for 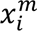. Thus, the ligand-anchor loss is defined as:

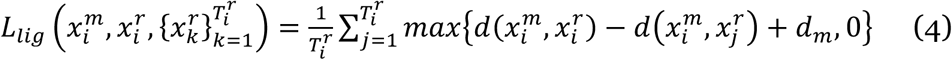

With the combination of the two symmetric loss terms, the final mini-batch loss for training the binding score module with batch size *N*_*b*_ is defined as

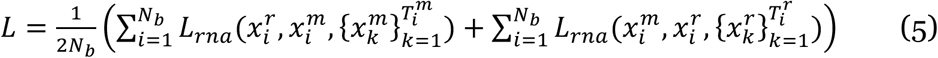

#### Ligand-specific decoy enhancement strategy

Although our negative sampling strategy enables the inclusion of all possible inactive binders in the training dataset for contrastive learning, current RNA-ligand interaction datasets, such as HARIBOSS, contain only a few hundred unique small molecules, representing a tiny fraction of available chemical space. The significant population differences of the decoy samples substantially hinder the model’s generalization ability, making it challenging to apply for practical virtual screening applications. To mitigate the differences and better reflect real-world scenarios, we propose a ligand-specific decoy enhancement strategy.

We first curated a large dataset of small molecules that are most unlikely to bind RNA targets, serving as a RNA-focused decoy library. Specifically, we used the bioactive molecules deposited in Inforna database^30^ and screened those against a subset of Enamine *REAL* database^31^ of more than 1 billion compounds. Both databases were converted to Morgan fingerprints and then compared using the Tversky similarity score^32^. Only candidate molecules with a Tversky similarity less than 0.4 to all Inforna compounds were retained. Compounds with a similarity score less than 0.4 were selected to create a library with approximately 2 million molecules. Then a two step clustering using the K-means approach was applied and the center of the clustres were selected to create a chemical diverse decoy library with 92,626 entries (Supplementary Note 2).

Instead of randomly sampling decoys from the curated library, we also considered molecular similarity to improve the decoy specificity corresponding to the active binders. We computed the pairwise Tanimoto similarity^33^ using FP2 fingerprints between each small molecules in the HARIBOSS dataset and compounds in our decoy library, and selected the top 200 most similar decoy per molecule. This process yielded an external, ligand-specific decoy set tailored to each active compound, substantially expanding the internal decoy set originally derived from the HARIBOSS dataset. During the model training, negative samples were drawn from both the HARIBOSS ligand pool and our curated external decoy library.

#### Attention-based RNA-ligand binding site prediction

In contrast to binding score prediction, which models global interactions between the RNA and a small moleucle, binding site prediction focuses on the nucleotide level, aiming to identify specific regions on the RNA sequence involved in ligand binding.

For the binding site prediction network, we employed an 8 multi-heads self-attention module with scaled dot-product attention and sinusoidal positional encoding to predict the binding site with the RNA sequence given its active binder. A layer normalization^81^ follows the attention to stabilize training and ensure consistent representations across sequence positions. The attention mechanism^88^ has shown its effectiveness in discerning intricate patterns critical for predictive models in biological sequences^89^. In this study, we first refine the representational capability of the RNA sequence embeddings generated by RNA-FM^25^ through the incorporation of sinusoidal positional encoding^88^. Subsequently, these enriched embeddings are concatenated with the small molecule embeddings at the nucleotide level. The concatenated embeddings are then processed through an MLP followed by batch normalization^90^ and a LeakyReLU activation^84^ function, to compress the combined features to a dimensionality that aligns with the small molecule embedding. A single-layer self-attention module with eight multi-attention heads is deployed to refine the feature representation, leveraging the inherent dependencies within the concatenated embeddings. This layer is followed by a residual connection^91^ that reintroduces the original small molecule representation. Finally, the refined features are passed through a binding site decoder network to predict the binding site.

Here, we employed a weighted binary cross-entropy (BCE) loss for training optimization. Specifically, a weighting factor is incorporated to address imbalanced classification during model training. It can be formulated as follows:

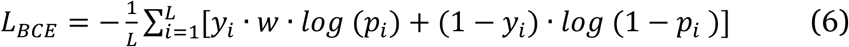

where *p*_*i*_ is the predicted probability that *i*-th RNA nucleotide is part of the small moleucle binding site, *y*_*i*_ ∈ {0,1} is the corresponding binary label, *w* is the weighting factor for positive residues (i.e., binding sites), and *L* is the RNA sequence length.

The Area Under the Receiver Operating Characteristic Curve (AUC) is used to evaluate the accuracy of binding site prediction. AUC scores were computed individually for each RNA–ligand pair in the test set, and the average score across all test samples was reported.

#### Ligand discovery using SMARTBind

To predict the interaction between an RNA sequence *r* and a small molecule *m*, the trained SMARTBind scoring function first generates the RNA embedding vector *x*^*r*^ and the small molecule embedding vector *x*^*m*^, mapping both inputs into the co-embedding space of dimenstion 256. The interaction likelihood is quantitatively assessed by computing the cosine similarity between these two embedding vectors. As one of the most commonly used methods for comparing embeddings in high dimensional spaces, cosine similarity ignores the vector magnitude and evaluates only orientation, making it scale-invariant and less sensitive to norm-related artifacts^27, 28, 92^. In our application, for a given RNA target, a higher calculated cosine similarity of a small molecule indicates a greater likelihood of the small molecule binding to it.

In the virtual screening scenario, our objective is to prioritize the most promising binder candidates, such as top-*k*, for an RNA target *r* from a large screening library of small molecules *M* = {*m*_1_, *m*_2_, …, *x*^*m*^}. We begin by calculating the embedding vector *x*^*r*^for the RNA and storing it in memory as the anchor for later comparison. We then compute the embedding vectors {*m*_1_, *m*_2_, …, *x*^*m*^} for all small molecules in the library and calculate their cosine similarities with the anchor vector. Based on these similarity scores, we rank the molecules and select the top-*k* candidates.

#### Methods for comparison

Few efforts have been investigated to the RNA-ligand interaction prediction field especially using data-driven appraoch. The closest peer-reviewed tools available during the development of SMARTBind are RNAmigos model series^34, 39^ and RNAsmol^46^. RNAmigos1^34^ represents 3D RNA binding pockets as augmented base-pairing graphs and employs graph neural networks to learn their embeddings, which are then matched to the Molecular Access Keys (MACCS) fingerprint^93^ of potential binders. RNAmigos2^39^ is the successor of RNAmigos1, incorporating directed edge-attributed RNA graphs, RNA language model-derived features, and deep graph encoders for ligands. Its training dataset was further expanded by integrating experimental data with docking-augmented data. For RNAmigos1 benchmarking, we adopted the reported results from their original publication and evaluated our model using the same validation dataset, ranking metrics, and experimental setting. For RNAmigos2, we obtained the mixed interaction score for each molecule against the query RNA pocket, for the benchmarking evaluation. Both methods require RNA binding pockets as input; therefore, we extracted the pockets using a 10 Å distance cutoff—consistent with their original configuration—when benchmarking on the HARIBOSS dataset and real-world case studies. Furthermore, RNAsmol^46^ is a supervised structure-agnostic deep learning framework that integrates data perturbation and augmentation, graph-based molecular representations, and attention-based feature fusion to predict RNA–ligand interactions without requiring structural inputs. We evaluated two model variants released by the authors: one trained on the PDB dataset, denoted as RNAsmol_PDB, and the other trained on the ROBIN dataset, referred to as RNAsmol_ROBIN.

In addition to the data-driven approach, we also benchmarked three widely used docking-based methods, including Dock 6^18^, rDock^40^, and AutoDock-GPU^4, 94^. DOCK 6 is an extension of DOCK 5 with improved sampling and scoring capabilities, optimization and testing for compatibility with RNA. rDock is designed to dock small molecules against proteins and nucleic acids. It employes a combination of stochastic and deterministic search algorithms to generate ligand binding poses and uses an empirical scoring function to evalute binding affinity. AutoDock-GPU is also a general-purpose docking program for both protein and RNA, based rapid Lamarckian genetic algorithm search method and an empirical free-energy force field.

#### Virtual screening against pre-miR-21

To design a structurally diverse and drug-like small molecule library for screening against pre-miR-21, we curated compounds from the ChEMBL database (∼2.5 million entries). We first applied a Quantitative Estimate of Drug-likeness (QED) filter, retaining only molecules with QED scores > 0.7, to enrich for compounds with favorable pharmacokinetic and physicochemical properties. The resulting subset was encoded using extended-connectivity fingerprints (ECFP4) and subjected to multiple rounds of K-means clustering based on Tanimoto similarity. To ensure both chemical diversity and practical manageability, the centroids (cluster centers) from each clustering iteration were selected as representative compounds for inclusion in the final library. This strategy yielded a curated set of structurally diverse, non-redundant, and experimentally tractable molecules. The resulting library was then employed for experimental screening against pre-miR-21, with the objective of identifying small molecules capable of binding or modulating its secondary structure or biogenesis.

#### Implementation details

##### Model training

Our model was implemented using PyTorch (version 2.0). The pre-trained RNA foundation model consists of 12 transformer layers, and only the last transformer layer was fine-tuned on RNA-ligand dataset while the other layers kept frozen to enable adaptation to the downstream RNA-ligand interaction task while preserving foundational RNA knowledge. All other SMARTBind modules were randomly initialized. The binding-score module was first trained using a contrastive loss over both positive and negative RNA–ligand pairs. Subsequently, the binding-site prediction module was integrated atop the frozen base model and trained exclusively on positive RNA–ligand pairs. Training was performed on a single Nvidia A100 GPU (80 GB). More details on model training and development are provided in Supplementary Note 4.

##### Docking methods benchmarking

To benchmark the performance of conventional docking approaches for RNA-targeted small molecule discovery, we evaluated Dock 6, AutoDock-GPU, and rDock on a curated dataset of 634 experimentally resolved RNA– small molecule complexes from the HARIBOSS database. For each complex, the native ligand was extracted, and the RNA target structure was preprocessed by removing non-relevant chains, water molecules, and ions. Ligand and receptor preparation followed standardized protocols: ligands were protonated and energy-minimized using Open Babel, and RNA structures were formatted using AutoDockTools (for AutoDock-GPU) and rbcavity/rdockprep (for rDock). Docking was performed using grid-based search spaces centered on the known ligand binding site, with box sizes defined to fully enclose the ligand-accessible pocket. Both rDock and AutoDock-GPU were run in a semi-blind docking mode, with the binding box spanning a broad region. This setup allowed for a fair comparison with our sequence-based model, which does not rely on predefined binding pockets.

We assessed scoring performance by calculating the rank percentile of the native ligand among all test set decoys, based on their top-1 poses generated by the docking algorithm. The rank percentile based on the docking scoring function provides a normalized measure of how well the native ligand score compares to the full set of decoys: a lower docking score corresponds to higher rank. These percentiles help evaluate whether the docking scoring function effectively prioritizes the biologically relevant poses. The analysis revealed systematic differences in pose prediction accuracy and rank-based prioritization between the three docking tools, highlighting the relative strengths and limitations of conventional scoring functions for RNA–ligand interactions.

##### RNA structure prediction

For RNA targets lacking experimentally resolved structures, we employed FARFAR2, a Rosetta-based fragment assembly protocol, to generate high-confidence 3D structural models. Starting from RNA sequences and predicted secondary structures (in dot-bracket notation), FARFAR2 assembles full-atom 3D conformations by sampling fragment libraries derived from known RNA motifs. Each modeling run produced an ensemble of candidate structures, from which the lowest-energy conformer was selected based on Rosetta all-atom scoring functions. Based on our experimentally validated reactive nucleotide A:19 for pre-miR-372, we extracted surrounding nucleotides within a 10 Å cutoff. The resulting pocket was then used as the input for RNAmigos1- and RNAmigos2-based virtual screening, enabling downstream evaluation of ligand binding against structurally uncharacterized RNA elements.

##### Ensemble virtual screening and binding site prediction

For the large-scale virtual screening case study and real-world miR-21 binder discovery presented in this study, we utilized the SMARTBind models trained with the 10-fold random split on HARIBOSS data, ensuring the model can utilize all the available RNA ligand information. An ensemble approach was employed for SMARTBind, in which binding score predictions were averaged across the ten cross-validation models to produce the final ranking output, while binding site predictions were similarly averaged across the ensemble and subsequently min–max normalized.

## DATA AVAILABILITY

RCSB PDB (https://www.rcsb.org/), HARIBOSS (https://hariboss.pasteur.cloud), and RNAmigos1 (https://github.com/cgoliver/RNAmigos) datasets are publicly available. SMARTBind resources and usage instructions are available via GitHub at https://github.com/AIDD-LiLab/SMARTBind under the MIT License.

## CODE AVAILABILITY

Source code for SMARTBind is publicly available via GitHub at https://github.com/AIDD-LiLab/SMARTBind under the MIT License.

## SUPPLEMENTARY INFORMATION

Supplementary Information includes Supplementary Notes explaining various features of the SMARTBind platform, Supplementary Figures 1 - 10, Supplementary Tables 1 – 3, and additional Experimental Methods.

