## Supplementary Information for "Small Molecule Approach to RNA Targeting Binder Discovery (SMARTBind) Using Deep Learning Without Structural Input"

### Contribute equally

#### SUPPLEMENTARY FIGURES & TABLES

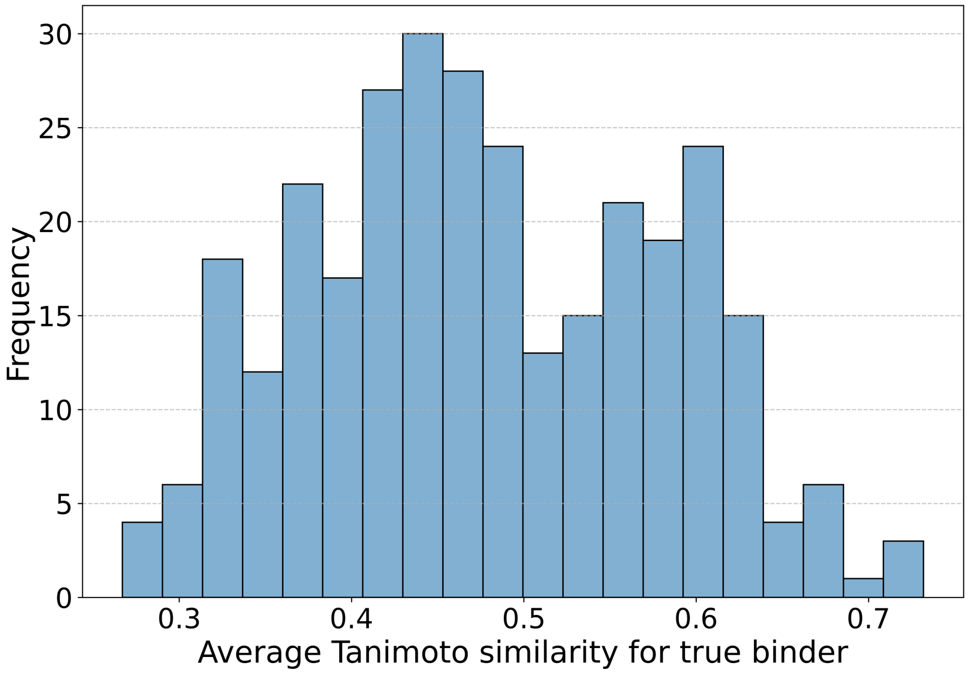


**Supplementary Fig. 1 | Distribution of Tanimoto similarity between ligand-specific decoys and their corresponding active binders.**

Histogram of average Tanimoto similarity calculated using FP2 fingerprint between each true binder and its 200 decoys selected by our ligand-specific decoy enhancement strategy. Most values fall between 0.4 and 0.6 (mean ± s.d.: 0.478 ± 0.101; median: 0.468), reflecting a balance between structural resemblance for challenging negatives with sufficient chemical diversity.

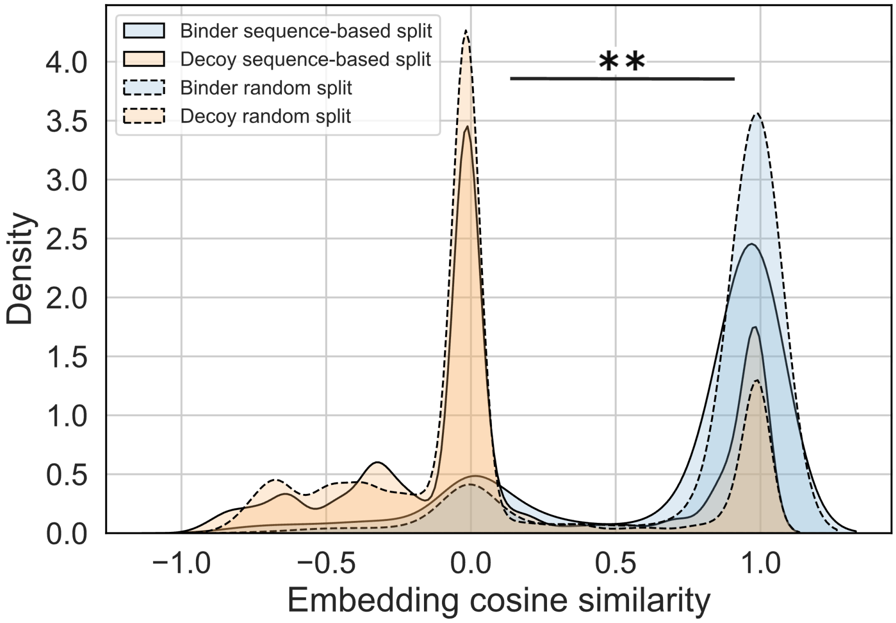


**Supplementary Fig. 2 | Embedding distance between binders and decoys in SMARTBind's latent space**

Density distributions of embedding cosine similarity scores between RNA sequences and their paired true binders or decoys, evaluated under sequence-based and random split conditions. A clear separation can be observed between the binder and decoy distributions across both settings, with true binders showing significantly higher embedding similarity to RNA sequences. Statistical significance was assessed using a one-sided Wilcoxon rank-sum test (** indicates $p < 0.01$, FDR corrected).

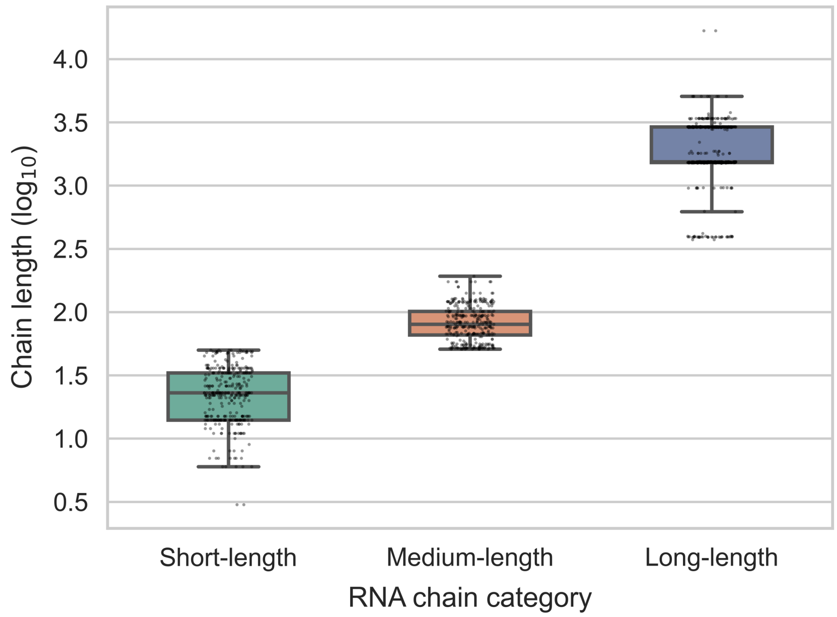


**Supplementary Fig. 3 | RNA chain length distribution across the HARIBOSS dataset.**

Box plots showing the RNA chain length distributions, displayed on a $\log_{10}$ scale, categorized as short (<50 nt; on average 24 ± 11 nt, mean ± s.d.; n = 359), medium (50-200 nt; on average 86 ± 28 nt; n = 345), and long (>200 nt; on average 2187 ± 1298 nt; n = 542) across the curated HARIBOSS dataset. Individual chains are shown as overlaid black dots.

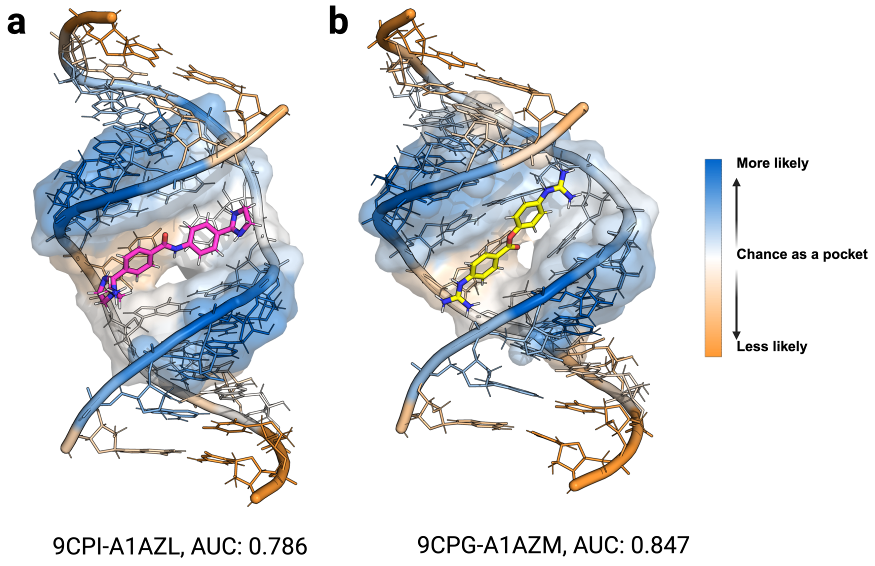


**Supplementary Fig. 4 | SMARTBind-predicted binding sites for active binders to the r(CUG) repeat RNA target.**

Predicted binding sites for active binders **(a)** A1AZL and **(b)** A1AZM to a r(CUG) repeat RNA (PDB IDs: 9CPI and 9CPG). Prediction confidence is visualized as a color gradient from blue (binding site, high confidence) to orange (non-binding site, low confidence). Surfaces indicate reference binding sites.

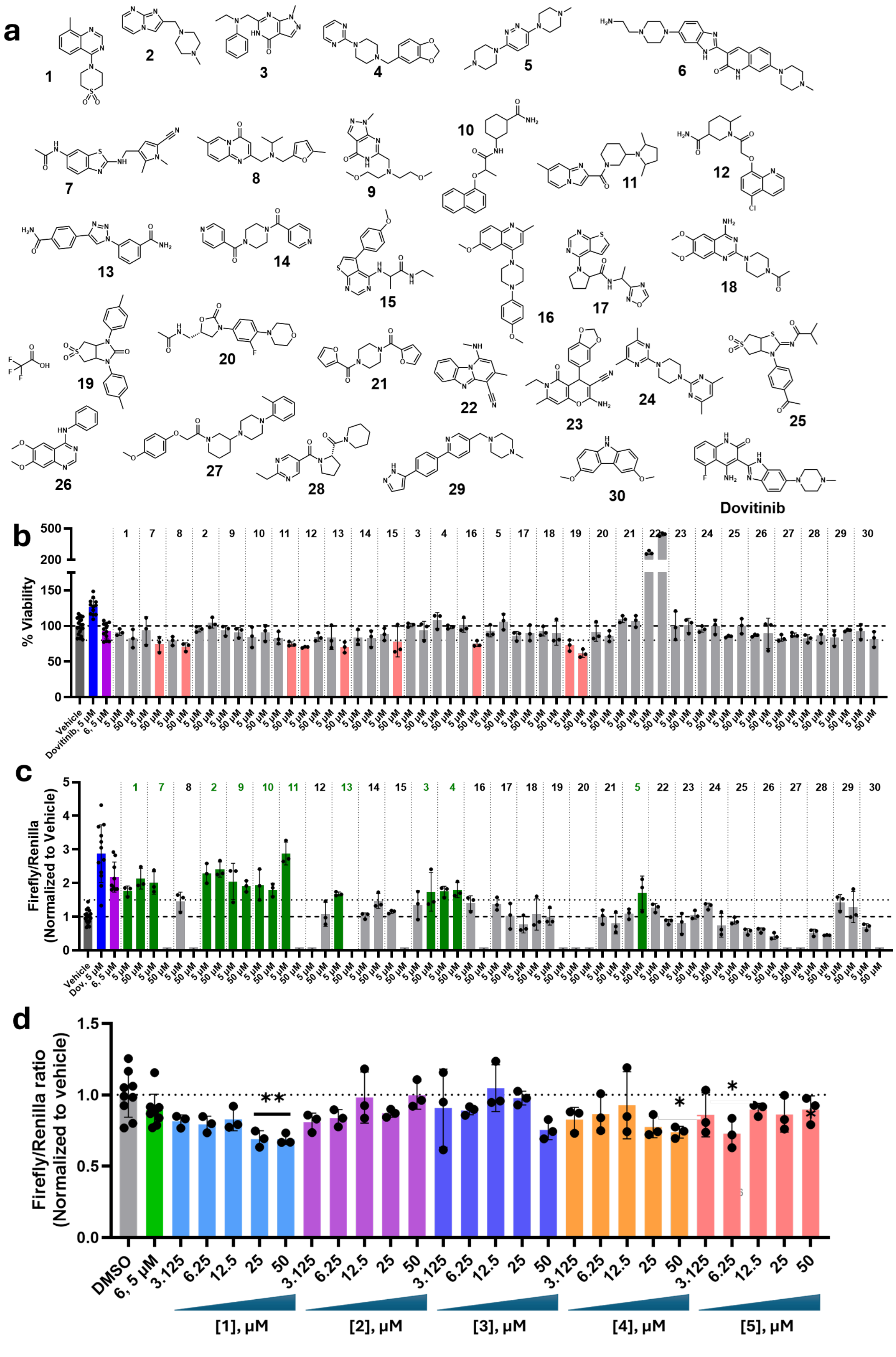

a
Dovitinib
b
% Viability
c
Firefly/Renilla (Normalized to Vehicle)
d
Firefly/Renilla ratio (Normalized to vehicle)
DMSO
6, 5 μM
[1], μM
[2], μM
[3], μM
[4], μM
[5], μM

**Supplementary Fig. 5 | Effects of 29 compounds designed by SMARTBind on cell viability and de-repression of PTEN in a luciferase reporter assay**

**(a)** Structures of the 29 SMARTBind-design compounds, dovitinib, and the dovitinib derivative **6**, the latter two of which serve as positive controls[1]. **(b)** Effect of the 29 compounds designed by SMARTBind, dovitinib, and **6** on the viability of MDA-MB-231 cells, as measured by Cell-Titer Fluor (n = 12 for vehicle, dovitinib, and **6**; n = 3 biological replicates for SMARTBind compounds). All 29 compounds were tolerated at 5 µM. Compounds 7, **8**, **11**, **15**, **16** and **20** were toxic (<75% viability) at the 50 µM dose. **(c)** Effect of the 29 compounds in a luciferase reporter assay where the 3' UTR of PTEN, a direct target of miR-21, is fused to firefly luciferase[2] (n = 12 for vehicle, dovitinib, and **6**; n = 3 biological replicates for SMARTBind compounds). Dovitinib and **6** were used as positive controls in the luciferase assay[1] . Compounds **1 – 5**, 7, **9-11**, and **13** increase luciferase activity to similar extent as **6** does (>1.5-fold increase). **(d)** Effect of **1 –5** in the PTEN luciferase assay conducted in HEK-293T cells (n = 9 for vehicle, dovitinib, and **6**; n = 3 biological replicates for SMARTBind compounds), which express miR-21 at much lower levels ($C_t$ = ~30) than MDA-MB-231 cells ($C_t$ = ~22; 256-fold higher expression)[3]. Thus, the compounds were expected to be inactive. **, $p < 0.01$; ***, $p < 0.001$; and ****, $p < 0.0001$, as determined by a one-way ANOVA with multiple comparisons. Data are reported as the mean ± SD.

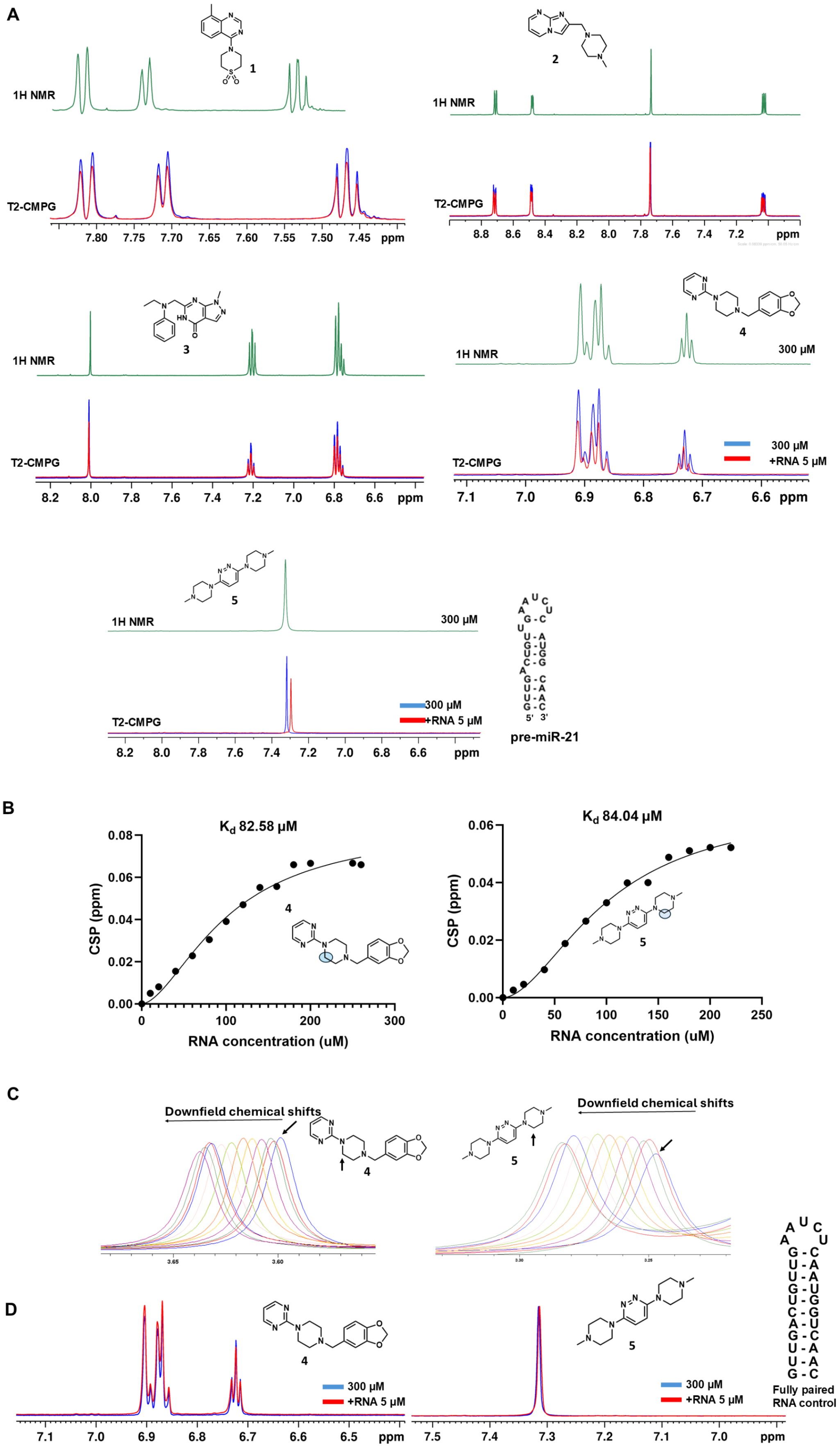
A
1
1H NMR
T2-CMPG
7.80 7.75 7.70 7.65 7.60 7.55 7.50 7.45 ppm
2
1H NMR
T2-CMPG
8.8 8.6 8.4 8.2 8.0 7.8 7.6 7.4 7.2 ppm
3
1H NMR
T2-CMPG
8.2 8.0 7.8 7.6 7.4 7.2 7.0 6.8 6.6 ppm
4
1H NMR
300 µM
T2-CMPG
300 µM
+RNA 5 µM
7.1 7.0 6.9 6.8 6.7 6.6 ppm
5
1H NMR
300 µM
T2-CMPG
300 µM
+RNA 5 µM
8.2 8.0 7.8 7.6 7.4 7.2 7.0 6.8 6.6 ppm
5' 3'
pre-miR-21
B
Kd 82.58 µM
CSP (ppm)
RNA concentration (uM)
4
Kd 84.04 µM
CSP (ppm)
RNA concentration (uM)
5
C
Downfield chemical shifts
4
3.65 3.60
Downfield chemical shifts
5
D
4
300 µM
+RNA 5 µM
7.1 7.0 6.9 6.8 6.7 6.6 6.5 ppm
5
300 µM
+RNA 5 µM
7.5 7.4 7.3 7.2 7.1 7.0 ppm
Fully paired RNA control

**Supplementary Fig. 6 | Compounds 4 and 5 bind specifically to the miR-21 Dicer site, as determined from T2-CPMG NMR studies.**

**(a)** CPMG spectra to investigate the binding of **1 – 5** to a model of miR-21's Dicer processing site. The proton spectrum for each compound alone is shown in green. Overlaid spectra in the absence and presence of the RNA are shown in blue and red, respectively. Broadened line shapes and reduced peak intensities are due to exchange between free and bound states after RNA addition. The difference spectra highlight attenuation arising from exchange-induced transverse relaxation in the presence of RNA, indicative of ligand binding. **(b)** 1D imino proton spectra of **4** and **5** as a function of concentration of the miR-21 Dicer site. The changes in chemical shift (chemical shift perturbation (CSP)) were plotted as a function of RNA concentration and fit to a specific binding model (GraphPad Prism) to afford the dissociation constant ($K_d$). **(c)** The corresponding proton peaks were used to calculate the $K_d$ for **4** and **5**, as indicated with black arrows. **(d)** T2-CPMG experiment with a fully paired RNA control, which showed no binding to **4** or **5**.

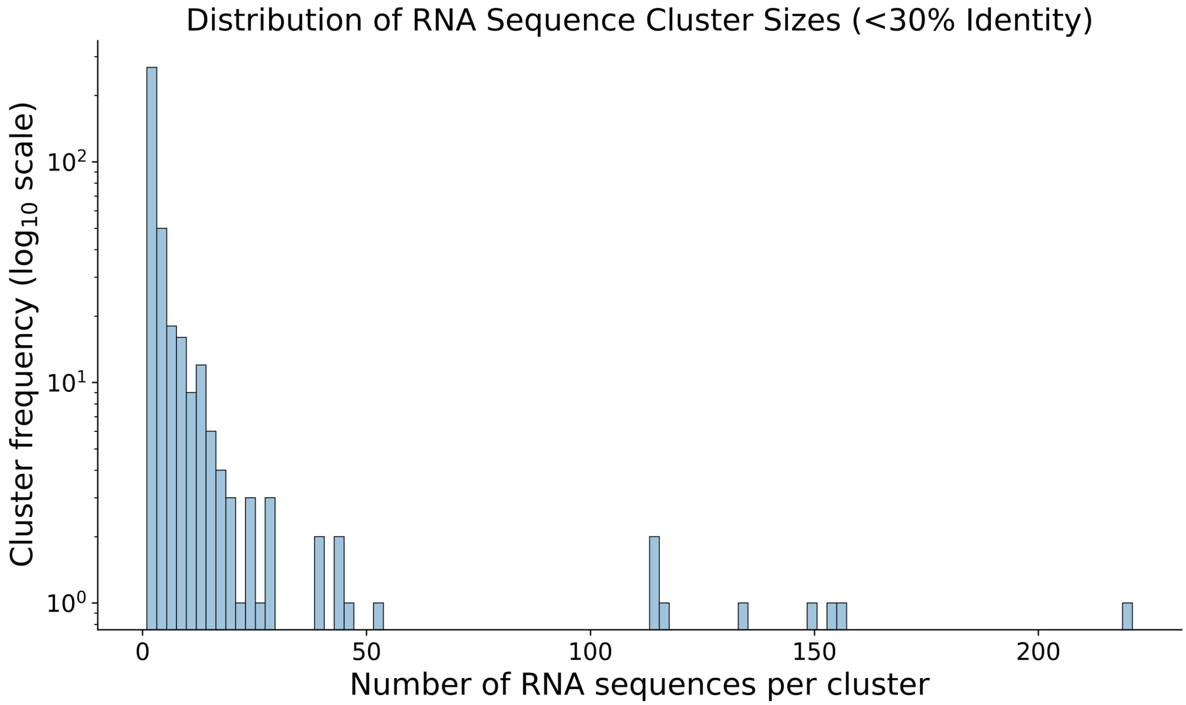


**Supplementary Fig. 7 | Distribution of sequence cluster sizes at <30% sequence identity.**

Histogram showing the frequency distribution ($log_{10}$ scale) of clusters constructed at 30% sequence similarity, plotted by the number of RNA sequences per cluster. The HARIBOSS dataset contains some RNA–ligand pairs with duplicated RNA sequences, resulting in a skewed cluster size distribution.

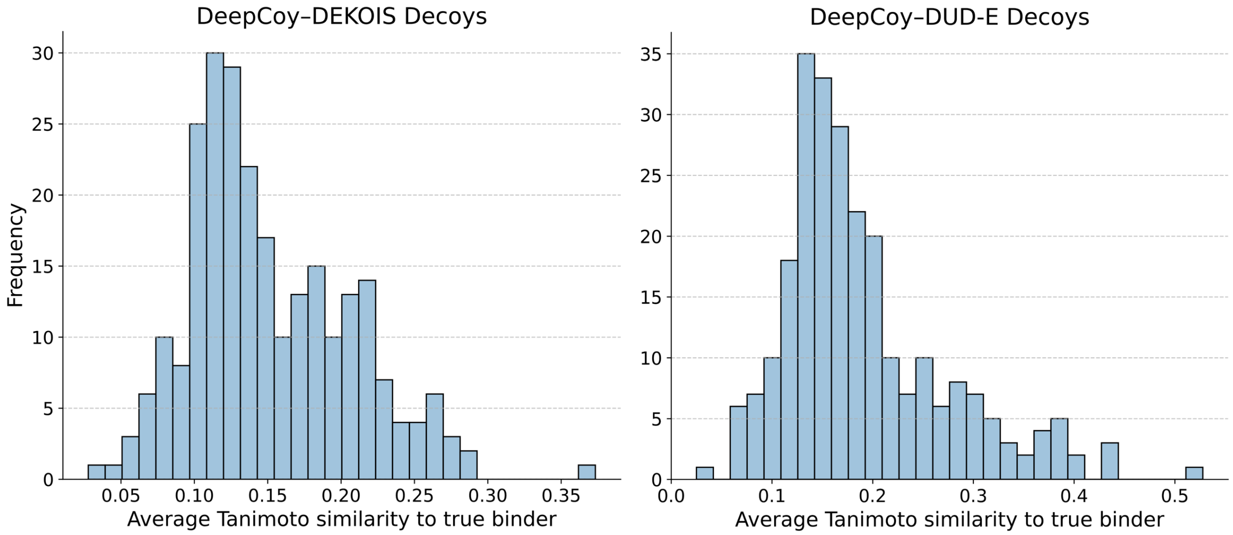


**Supplementary Fig. 8 | Distribution of Tanimoto similarity scores between DeepCoy generated decoys and their corresponding active binders.**

Histogram of average Tanimoto similarity calculated using FP2 fingerprint between each true binder and its 1,000 DeepCoy-generated decoys, using model variants trained on DEKOIS (mean ± s.d.: 0.152 ± 0.055; median: 0.139) and DUD-E (mean ± s.d.: 0.191 ± 0.082; median: 0.168) datasets.

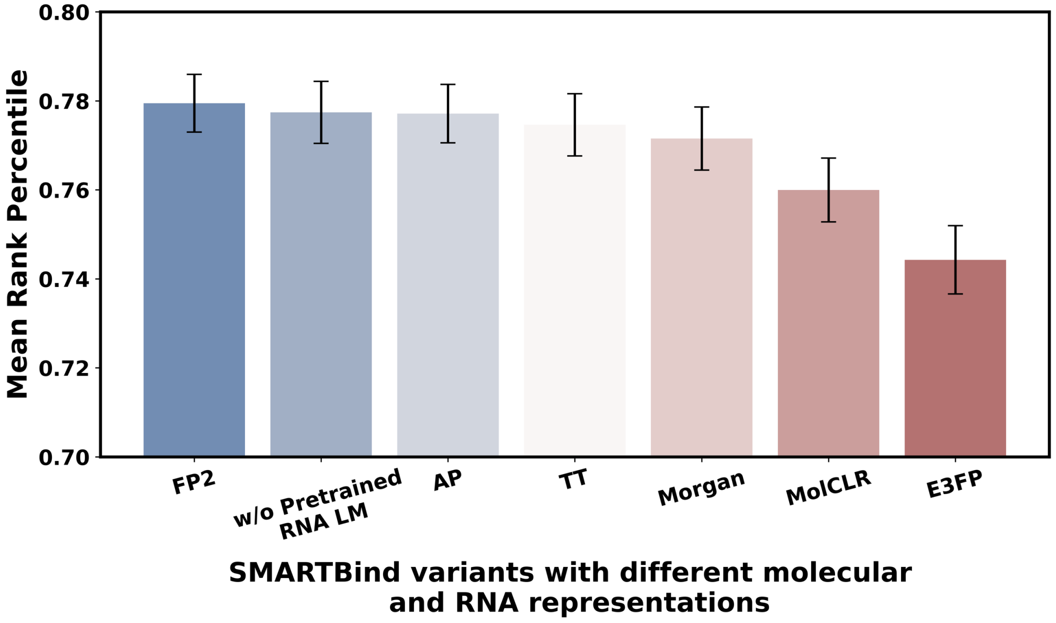


**Supplementary Fig. 9 | Performance comparison of SMARTBind with different molecular and RNA representations.**

Mean rank percentile performance comparation of SMARTBind ablation models using different molecular representations (FP2, AP, TT, Morgan, E3FP fingerprints, and MolCLR embedding), and with and without RNA language model pretraining. Error bars indicate the standard error of the mean (s.e.m.).

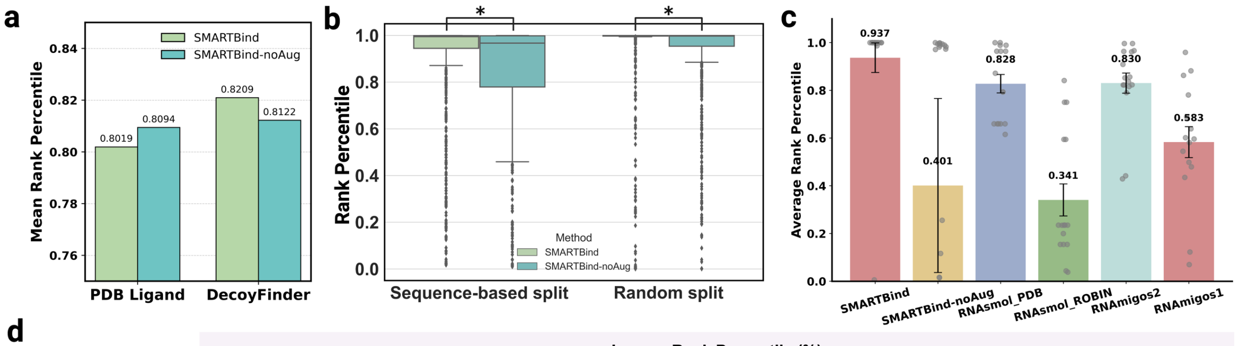


d

Inverse Rank Percentile (%)

| | Pseudoknot | | | G-quadruplex | Hairpin | | | Internal loop | | | | | | Groove platform | | N/A |
|---|---|---|---|---|---|---|---|---|---|---|---|---|---|---|---|---|
| | 8KED-GUN | 8KHH-GMP | 8KEB-GNG | 8K7W-2ZY | 8SYK-TAC | 8T5O-TAC | 8Z8Q-GNG | 8AZU-GUN | 9IOU-53D | 9IOS-53D | 9IO0-53D | 9IOR-53D | 9IO1-53D | 9MQT-A1BNU | 9FN2-SAH | 9MQS-5GP |
| SMARTBind | **0.0001** | **0.0019** | **0.0011** | **0.0218** | 1.31 | **0.24** | **0.0011** | **0.0001** | **0.0079** | **0.0066** | **0.0038** | **0.0011** | **0.0007** | **0.40** | 99.39 | 0.0004 |
| SMARTBind-noAug | 0.0003 | 0.5262 | 0.0891 | 1.07 | 98.51 | 74.41 | 0.1864 | 0.5363 | 2.87 | 0.9953 | 1.83 | 2.68 | 1.72 | 98.33 | 88.34 | 5.86 |
| RNAsmol_PDB | 20.65 | 3.62 | 3.62 | 38.46 | **1.06** | 1.10 | 3.61 | 20.64 | 34.02 | 34.01 | 34.01 | 34.00 | 34.01 | 12.80 | **0.0001** | **0.0001** |
| RNAsmol_ROBIN | 24.99 | 84.52 | 84.52 | 95.60 | 40.54 | 40.54 | 84.52 | 24.99 | 76.45 | 76.46 | 76.48 | 76.48 | 76.49 | 15.88 | 80.02 | 96.14 |
| RNAmigos1 | 92.97 | 21.91 | 36.08 | 57.09 | 13.71 | 4.05 | 31.07 | N/A | 39.79 | 40.37 | 50.12 | 87.71 | 52.04 | 56.47 | 11.82 | 41.88 |
| RNAmigos2 | 14.36 | 3.37 | 3.86 | 45.50 | 18.37 | 20.96 | 3.67 | 9.11 | 16.33 | 17.71 | 17.70 | 14.72 | 17.66 | 55.78 | 0.37 | 0.35 |

**Supplementary Fig. 10 | Performance comparison of SMARTBind and SMARTBind-noAug.**

**(a)** Performance comparison of SMARTBind and SMARTBind-noAug on the RNAmigos1-curated 10-fold random split dataset. Bar plots show mean rank percentiles for the PDB ligand- and DecoyFinder-based decoy evaluation sets. **(b)** Performance comparison of SMARTBind and SMARTBind-noAug on the HARIBOSS 5-fold sequence-based split dataset and 10-fold random split settings using a large-scale decoy evaluation set. The significance was assessed by a one-sided Wilcoxon rank-sum test (* indicates $p <$ 0.05, FDR-corrected). **(c)** Average rank percentile comparison for RNA–ligand complexes released in the PDB in 2025. Error bars represent standard error of the mean. **(d)** Inverse rank percentile performance for each target. Best results were highlighted in orange and shown in bold, the second best results are underlined.

**Supplementary Table 1. Temporally independent benchmark of RNA–ligand complexes released in 2025.** A curated test set of 16 RNA–ligand complexes released in the PDB in 2025 (collected on May 5, 2025). To prevent data leakage, RNA targets with sequences present in the training set (8VXZ, 8VY0, 8VY1, 8VXX) were excluded. Structural annotations for each RNA were retrieved from RNA3D Hub[4].

| PDB ID | Ligand Binder | Release Date | RNA Chain ID | 3D motif annotation |
|---|---|---|---|---|
| 8KED | GUN | 2025-02-19 | A | Pseudoknot |
| 8K7W | 2ZY | 2025-01-29 | A | G-quadruplex |
| 8KHH | GMP | 2025-02-26 | A | Pseudoknot |
| 8KEB | GNG | 2025-02-19 | A | Pseudoknot |
| 8SYK | TAC | 2025-02-19 | C | Hairpin |
| 8T5O | TAC | 2025-02-19 | A | Hairpin |
| 8ZAU | GUN | 2025-03-19 | A | Internal loop |
| 8Z8Q | GNG | 2025-03-19 | A | Hairpin |
| 9IOU | 53D | 2025-04-02 | A | Internal loop |
| 9IOS | 53D | 2025-04-02 | A | Internal loop |
| 9IO0 | 53D | 2025-04-02 | A | Internal loop |
| 9IOR | 53D | 2025-04-02 | A | Internal loop |
| 9IO1 | 53D | 2025-04-02 | A | Internal loop |
| 9MQT | A1BNU | 2025-04-30 | A | Minor groove platform |
| 9MQS | 5GP | 2025-04-30 | A | N/A |
| 9FN2 | SAH | 2025-01-22 | A | Major groove platform |

**Supplementary Table 2. Binding site prediction results for each RNA-ligand sample from the temporally independent benchmark using SMARTBind.**

| PDB ID | Ligand Binder | RNA Chain ID | AUC | RNA Length |
|---|---|---|---|---|
| 8KED | GUN | A | 0.907 | 72 |
| 8K7W | 2ZY | A | 0.804 | 49 |
| 8KHH | GMP | A | 0.936 | 71 |
| 8KEB | GNG | A | 0.905 | 72 |
| 8SYK | TAC | C | 0.803 | 107 |
| 8T5O | TAC | A | 0.525 | 124 |
| 8ZAU | GUN | A | 0.778 | 69 |
| 8Z8Q | GNG | A | 0.932 | 71 |
| 9MQT | A1BNU | A | 0.567 | 332 |
| 9MQS | 5GP | A | 0.636 | 332 |
| 9FN2 | SAH | A | 0.798 | 58 |
| 9IOU | 53D | A | 0.929 | 19 |
| 9IOS | 53D | A | 0.564 | 19 |
| 9IO0 | 53D | A | 0.718 | 19 |
| 9IOR | 53D | A | 0.625 | 19 |
| 9IO1 | 53D | A | 0.564 | 19 |

| **Supplementary Table 3. Sequences and catalog numbers for RT-qPCR primers used in this study.** | | |
|---|---|---|
| Primer name | Sequence or Catalog Number | Source |
| GAPDH fwd primer | TGCACCACCAACTGCTTAG | IDT |
| GAPDH rev primer | GATGCAGGGATGTTC | IDT |
| Pre-miR-21 fwd primer | CTGATGTTGACTGTTGAATC | IDT |
| Pre-miR-21 rev primer | GCCCATCGACTGGTGTTGCC | IDT |
| U6 TaqMan Primers | Assay ID 001973 | Thermo Fisher Scientific |
| miR-21 TaqMan Primers | Assay ID 000397 | Thermo Fisher Scientific |
| miR-21-targeting LNA oligonucleotide | Lot number 591028489 | IDT |

#### SUPPLEMENTARY NOTES

##### Supplementary Note 1: Ablation study of SMARTBind model

In this section, we evaluated various molecular fingerprinting strategies for representing small molecules within the SMARTBind-noAug framework, alongside an ablation analysis of the RNA foundation model. Each fingerprinting method offers distinct representational strengths and was assessed for its capacity to capture relevant chemical features in RNA-ligand interactions (Supplementary Figure 8). As described below, each fingerprinting method was used to encode small molecules into a 1024-dimensional one-hot embedding, followed a trainable, randomly initialized multi-layer perceptron (MLP) to map them into the RNA-ligand co-embedding space.

**FP2 Fingerprint** is a path-based fingerprint which indexes small molecule fragments based on linear segments of up to 7 atoms[5]. It captures molecular substructures through an enumeration of bond paths, resulting in a 1024-dimensional binary vector.

**Morgan Fingerprint**, also known as circular fingerprint, operates by iteratively generating a hash based on the connectivity of atoms up to a specified radius around each atom[6]. In our framework, we utilize the default radius, resulting in a 1024-dimensional vector that emphasizes the local molecular topology.

**Topological Torsion (TT) Fingerprint** encodes the molecular structure based on the torsion angle information between quadruples of bonded atoms, thus capturing stereochemistry and other three-dimensional aspects of molecular conformation[7]. It offers a distinct view from the path-based fingerprints.

**Atom Pair (AP) Fingerprint** catalogs pairs of atoms according to their types, separation (in terms of the number of bonds between them), and optionally, their relative stereochemistry[8]. It produces a count-based vector that can efficiently summarize long-range interactions within molecules.

**Extended 3-Dimensional Fingerprint (E3FP)** extends traditional 2D fingerprint concepts into three dimensions, capturing the spatial arrangement of atoms by folding distances between atom pairs into a fixed-length hash[9]. This method is particularly useful in modeling non-flat chemical entities and complexes, reflecting the 3D nature of molecular interactions.

We also evaluated the MolCLR[10] embedding for small molecule representation. MolCLR is a self-supervised learning framework trained on a large unlabeled dataset of approximately 10 million unique molecules using graph neural networks. This pre-trained model produces small molecule embeddings with a dimensionality of 512. Similarly, a MLP was introduced to map the MolCLR-derived small molecule representations. During RNA–ligand interaction learning, the weights of the MolCLR model were kept frozen to preserve its pre-trained knowledge.

For the comparison of small molecule representations, all other settings were kept constant and only the fingerprinting method was varied. The ablation study was conducted without ligand-specific augmentation strategy on the HARIBOS dataset with 5-fold sequence-based split. As shown in Supplementary Fig. 9, the model utilizing the FP2 fingerprint achieved better performance compared to alternative representations. Next, we evaluated the contribution of the pre-trained RNA language model by training the RNA encoder from scratch solely on the limited RNA–ligand dataset, with the FP2 fingerprint fixed as the small molecule representation for control. Without leveraging the pre-trained RNA foundation model, the model's performance dropped, highlighting the critical role of RNA knowledge transfer in enabling effective down-stream learning and improving predictive accuracy.

#### Supplementary Note 2: Construction of a decoy library using Inforna[11] and Enamine *REAL*

To construct a chemically diverse and RNA-relevant decoy library, we leveraged a query-based virtual screening approach using experimentally validated RNA-targeting small molecules housed in the Inforna database[11] and a large commercial compound collection (Enamine *REAL*). This strategy enables the systematic selection of decoy compounds that are topologically and chemically dissimilar to known RNA binders, providing a robust foundation for downstream benchmarking or classification model training.

The Inforna database was used as the source of validated RNA-binding small molecules[11]. Inforna is a curated repository of small molecules that bind structured RNA motifs, either from the literature or identified through two-dimensional combinatorial screening selection (2DCS) and structure–activity relationship (SAR) studies. All molecules that bind RNA secondary or tertiary structures were retrieved from Inforna and processed as SMILES strings.

The resulting query set consisted of 509 ligands (canonical SMILES), that bind to a diverse set of RNA structural targets including internal loops, hairpins, and G-quadruplexes. The Enamine *REAL* database (containing over 1 billion synthetically accessible compounds) was used as the search space. A filtered subset of this database containing ~2 million drug-like molecules (QED>0.7 MW 250–550 Da, logP $\leq$ 5, rotatable bonds $\leq$ 10) was used to reduce computational load while preserving chemical diversity. Molecular fingerprints were computed to encode chemical structures for both Enamine *REAL* and Inforna small molecules in a machine-readable format suitable for similarity comparison. Specifically, extended-connectivity fingerprints (ECFP4), also known as Morgan fingerprints (radius = 2), were calculated using RDKit.

For each Inforna query compound, the Tversky similarity index[12] was computed between the query and every candidate molecule in the filtered Enamine *REAL* subset. Two hyperparameters were set for the Tversky index, $\alpha$=0.85 and $\beta$=0.15. By adjusting these parameters, one can bias the similarity measure toward emphasizing features of one set over the other. Only candidate molecules with a Tversky similarity <0.4 to all Inforna compounds were retained. This threshold ensures the selected molecules are chemically dissimilar to known RNA binders and unlikely to exhibit RNA-binding activity.

To further enhance diversity and eliminate redundancy in the decoy set, the following filters were applied:

1. Duplicate SMILES and stereoisomers were removed.
2. Molecules with problematic substructures (e.g., pan-assay interference compounds, PAINS[13]) were filtered using SMARTS-based substructure filters[14].
3. Final decoy candidates were required to meet general drug-likeness criteria (e.g., Lipinski's rule of five compliance[15]).

All similarity calculations were performed using RDKit v2023.03.1, with batch fingerprint generation and Tversky scoring optimized using NumPy and multiprocessing for parallel efficiency. The entire workflow was executed on a Linux HPC cluster across 40 CPU cores.

To maximize chemical diversity in the final decoy set while avoiding overrepresentation of redundant scaffolds, we implemented a two-step clustering strategy:

- Initial Clustering of a Random Subset: A subset of the filtered decoy candidates (n = 1,6395,623) was clustered using k-means clustering on ECFP4 fingerprints[16].
- Centroid-Based Refinement of the Full Set: Cluster centroids from the first round were then used as reference points to re-cluster the entire filtered decoy pool. For each cluster, a single representative molecule closest to the centroid was selected, resulting in a final library of diverse and structurally representative decoys.

This two-tier approach avoids clustering biases due to large library size, enhances diversity coverage, and ensures robust representation of chemical subspaces that are dissimilar to RNA-binding ligands (n = 92,626).

#### Supplementary Note 3: Further comparison of SMARTBind and SMARTBind-noAug

We further evaluated SMARTBind against its non-augmented variant, SMARTBind-noAug, to quantify the contribution of the decoy enhancement strategy. As shown in Supplementary Fig. 10, two models achieved comparable performance on the RNAmigos1 dataset, with SMARTBind outperforming under the DecoyFinder decoy setting, but slightly underperforming under the PDB ligand setting.

Both methods were also compared on the HARIBOSS dataset under both 5-fold sequence-based split and 10-fold random split settings. To evaluate model robustness and generalization, a large-scale decoy evaluation set containing over 90,000 decoys for each true binder in the test set was constructed. These decoys were drawn from our curated decoy library of 92,626 entries, excluding molecules previously used for the ligand-specific decoy augmentation during the model training. As shown in Supplementary Fig. 10b, SMARTBind consistently outperformed SMARTBind-noAug across both split settings, achieving significantly higher rank percentiles ($p < 0.05$, FDR-corrected). These results demonstrate that our decoy enhancement strategy effectively improves SMARTBind's performance in binder discovery and reinforces its utility in realistic virtual screening scenarios.

A comparison in the large-scale virtual screening setting was also performed using the temporally independent benchmark comprising 16 RNA–ligand complexes released in the PDB in 2025 (collected on May 5, 2025). As shown in Supplementary Fig. 9c, SMARTBind (0.937 ± 0.062, mean ± s.e.m.) exhibits a marked improvement over SMARTBind-noAug (0.40 1 ± 0.364), highlighting the benefit of our decoy augmentation strategy. Meanwhile, SMARTBind-noAug ranked second in 11 of 16 cases across all six models (Supplementary Fig. 10d), indicating its effectiveness in more than half of the instances despite a lower average performance.

#### Supplementary Note 4: Model training and development details

For the contrastive co-embedding space, embeddings derived from RNA-FM[17] and MolCLR[10], and 1,024 one-hot representations from the fingerprints were processed using a dual-layer multilayer perceptron (MLP). Both the RNAs and the small molecules were mapped into a 256-dimensional space. The overall RNA representation was derived by employing mean pooling to aggregate nucleotide-level embeddings from RNA-FM. During training, only the final transformer layer of RNA-FM was fine-tuned to adapt the model to the RNA–ligand interaction task. SMARTBind model optimization was conducted using the AdamW optimizer[18] with default parameters, except for a weight decay of 0.001. Different learning rate schedulers were applied for the last layer of RNA-FM and other SMARTBind modules. For the former, a fixed learning rate of $1e^{-5}$ was maintained during the training. For the latter, cosine annealing with warm restarts[19] was employed, beginning with an initial learning rate of 0.01 and decaying to a minimum of 0.001 over eight iterations within each restart phase. A dropout ratio of 0.1 was applied across all MLP layers to mitigate overfitting[20]. Norm-based gradient clipping[21] was employed with a maximum gradient norm of 0.1. Training was performed in batches of 24 with early stopping based on validation loss monitoring. All hyperparameters were empirically determined based on benchmarking experiments.

To incorporate the ligand-specific decoy augmentation strategy into contrastive learning, 72 negative samples were generated for each true binder in a training batch by randomly selecting 48 decoys from the training ligand pool (excluding the true binders of the target RNA) and 24 decoys from the true-binder–specific decoy library, providing a balanced and diverse set of contrasts for representation learning.

For the binding site prediction task, an optimizer configuration consistent with the training above was employed. Dropout rates were set to 0.2 for the MLP and 0.4 for the self-attention layers. The binding site prediction module leverages frozen embeddings from the binding score prediction module for inference and optimization, with training incorporating an early stopping strategy to mitigate overfitting. The weighted binary cross-entropy (BCE) loss function was utilized, with a weighting factor of 12 to balance class distributions.

#### Supplementary Note 5: Molecular docking implementation

To prepare RNA-ligand complexes for structure-based docking, a Python-based preprocessing pipeline that processes mmCIF structures to extract relevant molecular components and generate docking-ready input files was developed. For each RNA-ligand complex, the pipeline performs the following steps. This automated pipeline enables the efficient preparation of large numbers of RNA-ligand complexes for virtual screening.

1. Model Selection and Filtering: The first model of each structure was extracted using Biopython's MMCIFParser. Non-essential components such as water molecules, ions (e.g., $Na^+$, $Cl^-$, $Mg^{2+}$), and small buffer additives (e.g., PEG, TRS) were removed using a custom residue whitelist filter.
2. Ligand and RNA Extraction: Ligands were identified by scanning for HETATM records labeled with the prefix H_. The ligand and the RNA (residues A, C, G, and U only) were extracted separately using PyMOL in headless mode. Each was saved as an individual mmCIF file.
3. Binding Pocket Definition: A spherical binding pocket was defined as all atoms within a 10 Å radius of the ligand. The selected atoms were extracted using a PyMOL script and saved in a separate file. This pocket was used to define the receptor region for docking.
4. Grid Parameter Calculation: The spatial coordinates of atoms in the binding pocket were used to calculate the grid center and box dimensions. These values were written to a .gpf file for downstream use with AutoDock grid map generation.
5. Format Conversion and Charge Assignment: Both the ligand and the receptor (binding pocket) were converted from mmCIF to PDBQT format using Open Babel (v3.1.1), with Gasteiger partial charges assigned.

6. Parallel Processing: All processing steps were executed in parallel using Python's multiprocessing module, allowing batch preparation across multiple CPU cores. Output files were organized into individual subdirectories based on the PDB ID.

To perform docking simulations, a companion Bash script that automatically generates and submits SLURM jobs for each RNA-ligand complex prepared in the previous step was developed. This docking pipeline streamlines the execution of GPU-accelerated AutoDock simulations across hundreds of RNA-ligand pairs with minimal manual intervention. The workflow proceeds as follows:

1. Input Parsing and Directory Matching: A CSV file listing complex IDs is provided as input. For each ID, the script performs a case-insensitive search to identify the corresponding directory containing the docking input files.
2. SLURM Job Generation: For each subdirectory, a SLURM job script is created. The job requests one GPU on the appropriate partition, loads required software modules, and sets the working directory to the subdirectory.
3. Docking Pipeline, within each SLURM job:
    a. prepare_gpf4.py and prepare_dpf4.py (AutoDockTools) are used to generate grid and docking parameter files, respectively.
    b. autogrid4 computes the receptor grid maps using the .gpf file.
    c. autodock_gpu_64wi (AutoDock-GPU v1.5.3) performs docking simulations using the precomputed grid maps and ligand PDBQT file.

4. Docked Pose Extraction: Docked conformations are extracted from the AutoDock.dlg log file by isolating lines starting with DOCKED. The coordinates are then converted to SDF format using Open Babel to facilitate downstream analysis.
5. Job Submission: Each SLURM script is submitted automatically using sbatch, enabling high-throughput docking on an HPC cluster.

**Software and Computational Environment:**

Biopython v1.79 – Structure parsing and manipulation

PyMOL v2.5+ – Structure selection and extraction in headless mode

Open Babel v3.1.1 – Format conversion and charge calculation

AutoDockTools v1.5.7 – Grid and docking parameter generation

AutoGrid v4.2.6 – Grid map calculation

AutoDock-GPU v1.5.3 – GPU-accelerated molecular docking

SLURM v20.11+ – Job scheduling on HPC

Python 3.8+ and Bash 5.0+ were used to execute the pipeline.

rDock-Based Docking of RNA-Ligand Complexes

To perform robust docking of RNA-ligand complexes using rDock, we implemented a fully automated pipeline in Bash. This script reads a list of RNA-ligand targets from a CSV file and performs a sequence of docking operations for each complex using rDock's receptor mapping and cavity generation workflow. An overview of the docking pipeline is as following:

1. Each row of the input CSV corresponds to a unique target, identified by a structure ID. For every target, the script performs the following steps:

- File Identification and Format Conversion: The script reads the receptor (*.cif) and ligand (*_igand.cif) files from a specified directory. These files are converted to rDock-compatible formats using Open Babel[22].
- Receptors are converted from mmCIF to MOL2 format and assigned Gasteiger partial charges.
- Ligands are converted from mmCIF to SDF format.

2. Docking Parameter Setup: A copy of the user-specified docking parameter file (dock.prm) is placed into the output directory of the current target. Additionally, a custom cavity parameter file (cavity.prm) is generated for each target. The cavity is defined using rDock's RbtLigandSiteMapper, centered on the reference ligand with a radius of 6.0 Å, and constrained to generate a single cavity.

3. Binding Site Grid Mapping: The cavity definition is computed using rbcavity[23], which generates the binding site grid file (binding_site.as) based on the previously defined cavity parameters.

4. Docking Execution: Ligands are docked using rbdock[23] with the generated cavity and docking parameter files. Each docking run performs 50 poses per ligand using rDock's built-in scoring function and the prepared binding site grid.

5. Post-processing and Pose Extraction: Upon successful docking, the generated .sd file containing all poses is renamed for consistency. The script then invokes a Python utility to extract the 25 lowest-energy docked poses and save them as a new SDF file.

Error Handling and Logging: The script includes robust error handling at each step. If required input files are missing, or if conversion, cavity generation, docking, or pose extraction fails, informative messages are printed and the pipeline continues to the next target.

**Software and Computational Environment:**

rDock v0.5.1 – Ligand docking, cavity detection, and scoring

Open Babel v3.1.1 – Molecular format conversion and charge calculation

CUDA v11.4, GCC v9.3.0, OpenMPI v4.0.5, Amber tools Python 3.8+ – Used for post-docking pose extraction

The pipeline is designed for execution in a SLURM-based HPC environment with appropriate module loading.

**Input and Output Structure**

Input CSV: A single-column CSV file listing structure IDs (case-insensitive).

Base Directory: Contains one subdirectory per structure, with receptor and ligand files named consistently (*_rna.cif, *_ligand.cif).

Outputs per target: Converted receptor and ligand files (.mol2, .sdf), docking and cavity parameter files, binding site grid file (binding_site.as), full docking output (docking_results.sdf), top 25 low-energy poses (lowest_energy_poses.sdf).

**Execution Summary**

For each target in CSV:

→ Convert RNA receptor (CIF → MOL2)

→ Convert ligand (CIF → SDF)

→ Generate docking and cavity parameter files

→ Run rbcavity to define binding site grid

→ Execute rDock docking for 50 poses

→ Extract and save 25 lowest-energy poses

This modular design supports rapid docking of large ligand libraries across multiple RNA targets and integrates seamlessly with upstream preprocessing and downstream hit prioritization pipelines.

#### MATERIALS & METHODS

**Small molecules:** The 29 SmartBind-designed small molecules were obtained from ChemSpace. The purity of the compounds was evaluated by high performance liquid chromatography while identity was confirmed by high resolution mass spectrometry.

##### Cellular Methods

**Cell culture.** MDA-MB-231 cells (ATCC, catalog #HTB-26,) were cultured in Roswell Park Memorial Institute Medium (RPMI; Corning, catalog #MT10041CV) supplemented with 10% (v/v) fetal bovine serum (FBS; Sigma Aldrich, catalog # A36707-01) and 1% penicillin-streptomycin (PS; Corning, catalog #30-002-CI). Cells were maintained at 37˚C and 5% $CO_2$. Upon reaching ~40% confluency, cells were treated by adding fresh growth medium containing the compound of interest, diluted from 1000× compound stocks (final DMSO concentration 0.1% (v/v)). After 24 h, the compound-containing growth medium was replaced (re-dosing) for a total treatment time of 48 h unless otherwise noted. The passage number of MDA-MB-231 cells used in all experiments was <20.

HEK293T cells (ATCC, catalog #CRL-3216) were maintained in Dulbecco's Modified Eagle Medium (DMEM; Corning, catalog 15-017-CV) supplemented with 10% (v/v) fetal bovine serum (FBS; Sigma Aldrich , catalog #A36707-01), 1% (v/v) Penicillin-Streptomycin Solution (Corning, catalog #30-002-CI) and 1% glutaGRO supplement (Corning, catalog #25-015-CI) at 37°C and 5% CO2. The passage number of HEK-2935 cells used in all experiments was <20. Both cell lines were tested for mycoplasma contamination, which was negative.

**PTEN luciferase assay and cell viability.** MDA-MB-231 cells were grown in 60 mm diameter dishes until they reached ~80% confluency. Cells were then transfected with 4 µg of a plasmid encoding the PTEN 3' UTR fused to firefly luciferase (Addgene # 21327)[3] and 0.8 µg of a Renilla-encoding plasmid (used for normalization) using jetPRIME transfection reagent (Polyplus) per the manufacturer's instructions for forward transfection. Approximately 6 h post-transfection, the cells were harvested and countered using Invitrogen Countess II Automated Cell Counter, seeded into 96-well plates (10,000 cells / well), and allowed to adhere overnight. The cells were then treated with compound prepared in grown medium, as described in "Cell Culture." Following the 48 h-treatment period, cell viability was measured using CellTiter-Fluor™ Cell Viability Assay (Promega) per the manufacturer's instructions. Fluorescence was measured using a Molecular Devices SpectraMax M5 plate reader. Luminescence was then measured in the same well using a Dual Glo Luciferase Assay System (Promega, catalog # E2920) per the manufacturer's protocol using a Tecan Infinite M1000 Pro plate reader. Data are reported as the ratio of LucFirefly (PTEN) to LucRenilla (internal control), normalized to vehicle-treated samples. HEK-293T cells were transfected and tested following the same protocol.

**Reverse transcription-real-time quantitative polymerase chain reaction (RT-qPCR) to measure pre-miR-21 and miR-21-3p abundance.** MDA-MB-231 cells were cultured in 12-well plates until they reached ~30% confluency. They were then treated with small molecule as described in "Cell Culture." After treatment for 48 h, total RNA was extracted from the cells using a Quick-RNA Miniprep Kit (Zymo Research), according to the manufacturer's instructions, including the on-column DNase I digestion. Total RNA was quantified by its absorbance at 260 nm using a Nanodrop

UV/Vis spectrophotometer (Thermo Fisher); only samples with $OD_{260}/OD_{280}$ >1.8, were further processed.

For measurement of pre-miR-21 abundance, 200 ng of total RNA was reverse transcribed using the qScript™ cDNA Synthesis Kit (Quantabio), per the manufacturer's protocol in a total volume of 20µl. RT-qPCR samples contained 1× Power SYBR Green Master Mix (Applied Biosystems), 25 ng of cDNA generated from the qScript™ RT reaction, and designed SYBR primers (57 nM each; see Supplementary Table 2 for sequences) in a total volume of 35 µL. The sample was then aliquoted into three 10 µL technical replicates. Amplification was completed using a QuantStudio5TM Real-Time PCR Instrument (Applied Biosystems).

For mature miR-21 abundance measurement, 100 ng of total RNA was reverse transcribed using a TaqMan MicroRNA Reverse Transcription Kit (Thermo Fisher) per the manufacturer's protocol in a total volume of 15µl. The corresponding qPCR samples were prepared using a TaqMan Fast Advanced Master Mix (Thermo Fisher), 25 ng of cDNA generated from the TaqMan RT reaction, and TaqMan primers (Supplementary Table 2). Amplification was completed using a QuantStudio5TM Real-Time PCR Instrument.

All RT-qPCR data were analyzed using the ΔΔCt method[24]. Expression levels of pre-miR-21 were normalized to GAPDH while expression levels of miR-21 were normalized to U6 snRNA.

**Simple Western to measure PDCD4 abundance**. MDA-MB-231 cells were cultured in 6-well plates until they reached ~30% confluency. They were then treated as described in "Cell Culture." After the 48 h-treatment period, the medium was removed, and the cells were washed with 1× DPBS. After removing the 1× DPBS, total protein was extracted using 200 μL of Mammalian Protein Extraction Reagent (MPER; ThermoScientific) with 1× Protease Inhibitor Cocktail III (RPI P50700-1) per the manufacturer's protocol. Protein concentration was quantified by using a Pierce BCA kit (catalog #23255) in 96-well plate per manufacturer's protocol. To measure endogenous PDCD4 protein, a Simple western was completed using a Jess Automated Western Blot System (Protein Simple) per the manufacturer's protocol. The concentration of protein loaded onto the Simple Western was 0.4 mg/mL. The β-Actin primary antibody (Cell Signaling Technology, catalog #8H10D10) and PDCD4 primary antibody (Cell Signaling Technology, catalog #D29C6) were diluted 1:100. Chemiluminescence mode and 25 capillary cartridges were used. Data are reported as the ratio of PDCD4 to β-Actin, normalized to vehicle-treated samples.

**Caspase-3/7 activity**. MDA-MB-231 cells were cultured in 96-well plates until they reached ~30% confluency. They were then treated as described in "Cell Culture." After 48 h treatment, Caspase-3/7 activity was measured using Caspase-Glo® 3/7 Assay System (Promega, catalog #G8090) per the manufacturer's recommended protocol. Luminescence was measured using a Tecan Infinite M1000 Pro plate reader (integration time =500 ms) Data is normalized to vehicle-treated samples.

#### In vitro Methods

##### NMR Spectroscopy.

NMR spectra for Carr–Purcell–Meiboom–Gill (CPMG)[25] and 1D imino and aromatic proton spectra were acquired on a Bruker Advance III 600 MHz spectrometer equipped with a cryoprobe. All spectra for were processed using TopSpin 4.1.1 (Bruker).

The model of the pre-miR-21 Dicer site used in NMR studies was previously reported[1] and has the following sequence: 5'-GUGUUGACUGUUGAAUCUCAUGG_CAACAC-3' where the A bulge and the missing cross-strand nucleotide are underlined (purchased from Dharmacon as an HPLC-purified and desalted oligonucleotide). The sequence of the fully base-paired RNA control was: 5'-GUGGACUGUUGAAUCUCAAUGGUCAAAC-3'. All nucleotides are endogenous to pre-miR-21. The RNA was diluted in NMR Buffer (10 mM $Na_2HPO_4/NaH_2PO_4$, pH 6.0, and 0.05 mM EDTA) and folded by heating at 95 °C for 3 min followed by snap cooling on ice.

Samples for CPMG experiments contained 5% (v/v) $D_2O$ (Cambridge Isotope Labs), 300 μM compound, and 5 μM RNA in a final volume of 600 μL. Experiments were carried out by first collecting spectra of **1** – **5** alone at a concentration of 300 μM, followed by addition of folded pre-miR-21 Dicer site (5 μM), affording a final ratio of RNA/compound of 60:1. To quantify transverse relaxation times ($T_2$) of RNA-ligand complexes, a pseudo-2D CPMG pulse sequence combined with excitation sculpting-based water suppression, implemented as the cpmg_esgp2d program on a Bruker Avance NMR spectrometer was employed. This sequence enables reliable detection of spin echo decay while effectively suppressing the water signal.

The experiment was configured such that a series of 1D experiments were recorded with incrementally increasing numbers of spin echoes using the default settings (cpmglist). The echo spacing (d20) was chosen to be significantly shorter than 1/J but long enough to accommodate the 180° refocusing pulses, ensuring well-resolved echo formation. The number of echoes in each train (controlled by COUNTER1) was incremented across the indirect dimension (td1), enabling exponential $T_2$ decay curves to be reconstructed for each signal.

Water suppression was achieved via excitation sculpting, which uses a pair of shaped 180° pulses bracketed by pulsed field gradients (gp1, gp2). This approach selectively inverts and refocuses water magnetization, while leaving solute signals unaffected. Phase cycling schemes were applied to suppress artifacts and select desired coherence pathways.

To quantify the binding affinity of **4** and **5**, chemical shift perturbations (CSP) of protons in **4** and **5** protons was measured as a function of miR-21 Dicer site concentration. A 50 μM solution of **4** or **5** were prepared in a final volume of 400 μL (Shigemi NMR tube). RNA (0.5 μL aliquots) was then added to afford concentrations ranging from 10-250 μM. The resulting curves plotting changes in CSPs as a function of RNA concentration were fitted using the one site specific binding model within GraphPad Prism.
